# Germinal center BCR maturation in appendicitis reveals a role for antigen-specific adaptive immune responses during disease

**DOI:** 10.1101/2024.01.31.578312

**Authors:** Emma Stewart, Zainab Taghvaei, Carmen Leon, Jason Shapiro, Lisha Zhu, Lindsay Alpert, Kiran Turaga, Roshni Roy Chowdhury

## Abstract

Appendicitis is one of the most common abdominal emergencies globally, yet little is understood about the inflammatory mechanisms or potential drivers of disease. Neutrophil inflammation and increased cytokine expression such as IL-6 and IL-8 are hallmarks of appendicitis inflammation. However, early histological studies identified increased T and B cell infiltration during appendicitis, providing support for adaptive immune activation as well, although this has never been investigated in depth. We hypothesized that antigen-dependent activation of the adaptive immune response contributes to appendicitis pathology, in addition to the known innate-mediated processes. Via a series of transcriptomic approaches and lymphocyte repertoire analysis in human appendiceal tissue, we identified evidence of antigen-dependent B cell activation. Increased somatic hypermutation in the germinal center and plasma cell compartment was comprised of presumed high-affinity IgG and IgA B cells. We propose that the appendiceal microbiome acts as a source of antigen, as significant microbial dysbiosis was observed during appendicitis. This dysbiosis was characterized by outgrowth of pathobionts such as *Parvimonas* and oral biofilm-formers such as *Fretibacterium* and *Fusobacterium*, in line with previous reports. We also identified potential loss of epithelial barrier integrity via spatial transcriptomic analysis of the appendiceal epithelium, supporting the possibility of microbial invasion into the tissue during appendicitis. This study provides insight into the inflammatory mechanisms of a common disease and helps to define the immune and microbial compartment of an often-ignored organ, the appendix.

## Introduction

Appendicitis is one of the most common abdominal emergencies, with an estimated prevalence of 8.7% globally^1^. Historically, appendicitis was thought to be caused by obstruction of the appendiceal orifice via fecalith or tumor, although current evidence suggests this is the exception rather than the rule, and cannot explain the majority of cases^2^. Microbial dysbiosis in the appendix, host genetics, environmental changes, or infection have all been proposed as causative factors, however, neither the precise etiology of appendicitis nor the molecular basis of the inflammatory response are well understood.

Transmural neutrophil infiltration is the hallmark feature of acute appendicitis. However, early histological studies have shown elevated T cell and plasma cell infiltration in appendicitis tissue compared to normal appendices^3^. Yet, a systematic analysis of the adaptive immune response in appendicitis is absent from the literature. A recent study has identified positive correlations between B cell infiltration into the appendix with abundance of certain microbial genera^4^. The association between the microbiome and immune response in appendicitis is reasonable, as the appendix is both a site of dense microbial life and lymphoid tissue^3, 5^. Multiple groups have investigated the microbiome during appendicitis. From these studies a few common findings have emerged, including a relative decrease in *Bacteroides* and increases in oral pathobionts such as *Fusobacterium, Gemella* and *Porphyromonas* in appendicitis samples^6–8^. The degree of dysbiosis appears to correlate with disease severity, with oral taxa and opportunistic pathogens such as *Parvimonas micra* more often enriched in complicated appendicitis characterized by gangrenous or perforated tissue, while uncomplicated cases had a lesser dysbiosis composed of typical gut commensal taxa^9–11^. Further implicating the microbiome in disease etiology is the result of the APPAC randomized clinical trial, which demonstrated that antibiotic therapy alone can be effective in some cases of appendicitis^12^. However, use of antibiotics alone has been shown to have a 25- 40% failure rate in long term follow up studies, with inflammation recurring up to years after the initial incident^13, 14^. This suggests a more complicated disease course than can be explained by bacterial infection alone. Understanding the underlying disease mechanisms in appendicitis is crucial for optimization of treatment strategies.

Thus far, research on the role of the immune response in appendicitis has focused on innate immunity. A past study found a polymorphism resulting in decreased IL-6 production was associated with reduced risk of complicated appendicitis^15^. An observational study found serum levels of IL-6 and IL-10 were increased at the time of appendicitis compared to one month later, and serum IL-6, IL-8, and TNF-α were all increased in complicated versus uncomplicated disease^16^. To our knowledge, no one has yet investigated the role of adaptive immunity, namely T and B cells, and their contribution to appendicitis pathology. While the microbiota and bacterial infiltration into appendiceal tissue have been proposed as a contributing factor, there is also no knowledge of bacterial antigens that may contribute to appendicitis inflammation.

Using a series of transcriptomic approaches, we profiled the gene expression profiles of appendiceal immune cells, TCR and BCR repertoires, and microbial communities from appendicitis samples. In the course of this analysis, we identified expanded IgG+ and IgA+ B cell clonotypes present in the germinal center and plasma cell compartments with increased mutational frequency. This finding indicates antigen-dependent and T cell-dependent BCR activation and maturation occurs during appendicitis. Antigen-driven T cell activation was less obvious, although Tfh cells that support B cell activation in the germinal center were identified. Analysis of the appendiceal microbiome during appendicitis revealed alterations in the community composition, particularly in perforated appendicitis, with obvious outgrowth of *Parvimonas*. We also identified a network of bacterial genera that were positively correlated with one another in appendicitis which included *Parvimonas, Peptostreptococcus, Fusobacterium,* and *Fretibacterium*. We hypothesized that loss of epithelial barrier integrity as well as invasive capabilities of bacterial species could lead to tissue invasion of microbes into the appendix, and confirmed gene signatures in the appendiceal epithelium were consistent with barrier stress. Finally, we wanted to identify potential antigens derived from the appendicitis microbiome. As BCR epitopes are technically challenging to predict, we utilized computational prediction tool to identify MHCII restricted peptides that could potentially be recognized by CD4+ Tfh cells as they contribute to B cell responses in the follicle. Overall, this analysis advances our understanding of the adaptive immune response during appendicitis, as well as the potential role of the microbiota in contributing to disease pathology.

## Results

In order to explore the immune response in appendicitis, we collected archival formalin-fixed paraffin embedded (FFPE) tissue sections from three appendicitis cases who underwent appendectomy. Four age-matched normal appendix tissue sections were also collected from patients undergoing incidental appendectomy. Demographic information for samples used in the spatial analysis can be found in **Table 1**. We performed exploratory spatial transcriptomic analysis using the Nanostring GeoMx platform. Sections were stained to highlight B cells (CD20), T cells (CD3), and epithelial cells (pan-cytokeratin) to identify regions of interest (ROIs) for the spatial analysis. ROIs containing greater than 300 estimated cells were collected across appendicitis and normal donors for 2 compartments: (1) Follicular B cells (“FB”) and (2) Follicle-adjacent T cells (“FT) (**Figure 1A**). After data normalization 8,805 genes were identified for downstream analysis.

**Figure 1.**
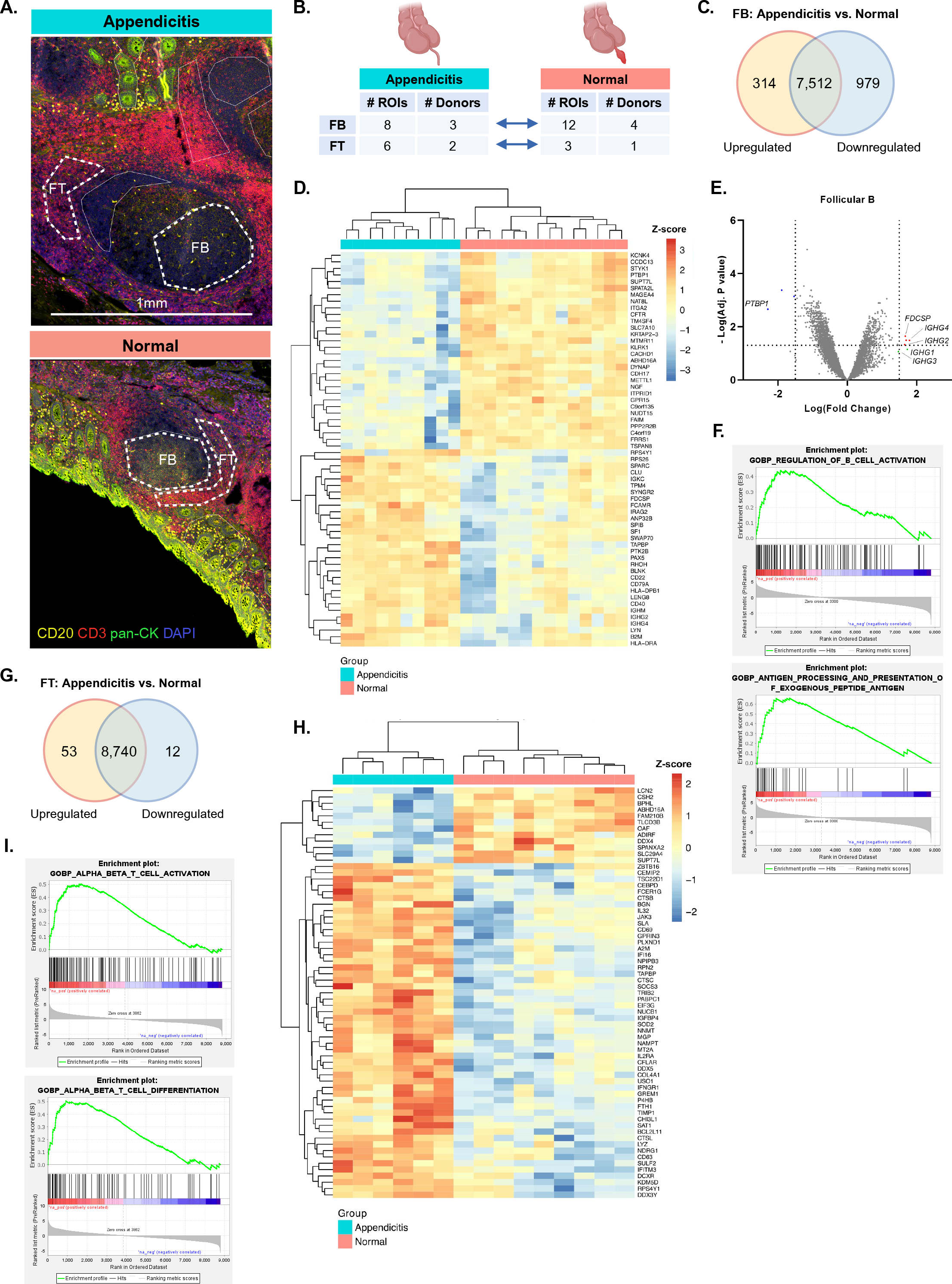
Appendiceal lymphocyte follicles show transcriptional upregulation of T and B cell activation in appendicitis, supporting a role for the adaptive immune response during disease. **(A)** Representative ROIs used in spatial transcriptomic analysis appendiceal FFPE sections. ROIs are outlined with a white dashed line and labelled “FB” or “FT”, sections are colored for CD20 (yellow), CD3 (red), pan-cytokeratin (green), DAPI (blue) and labelled with 1mm scale bar. **(B)** workflow schema for spatial transcriptomic analysis summarizing number of ROIs and donors for each comparison. **(C)** Summary of DE genes from FB analysis, based on adjusted p-value cutoff of ≤ 0.05. **(D)** Heatmap showing the top 30 upregulated and downregulated DE genes from FB segments. Heatmap is colored by Z-score of gene expression, with induvial genes labelled along the right side of the plot. **(E)** Volcano plot for DE genes from FB segments, horizontal dashed line indicates adjusted p-value cutoff of ≤ 0.05, while vertical dashed line indicates log(FC) ≥ ± 1.5. Genes upregulated in appendicitis are colored red, while downregulated genes are colored in blue. Relevant significant genes are annotated. **(F)** Representative GSEA plots showing enriched B cell activation and antigen processing and presentation from the FB segments. GSEA pathway significance was determined using and FWER p-value cutoff of ≤ 0.05. **(G)** Summary of DE genes from FT analysis, based on adjusted p-value cutoff of ≤ 0.05 **(H)** Heatmap showing all significant DE genes from FT segments, colored by Z-score of gene expression with gene labels along the right. **(I)** Representative GSEA plots showing enrichment in T cell activation and differentiation pathways from the FT segments. GSEA pathway significance was determined using an FWER p-value cutoff of ≤ 0.05. ***Abbreviations:*** ROI = region of interest, FB = follicular B regions, FT = follicle-adjacent T regions, FFPE = formalin fixed paraffin embedded, DE = differentially expressed, FC = fold change, GSEA = gene set enrichment analysis, FWER = family-wise error rate

**Table 1.** Spatial transcriptomic cohort demographic and clinical characteristics.

| Donor ID | Study Group | Severity | Diagnosis/Indication | Surgery | Age | Sex | Race |
| --- | --- | --- | --- | --- | --- | --- | --- |
| N1 | Normal | NA | Ovarian adenocarcinoma | Cytoreduction | 30 | F | Unknown |
| N2 | Normal | NA | Benign ovarian cyst | Appendectomy | 33 | F | Black |
| N3 | Normal | NA | Ovarian cystadenoma | Salpingo-oophorectomy, omentectomy, appendectomy | 38 | F | Unknown |
| N4 | Normal | NA | Gunshot wound | Appendectomy | 21 | M | Black |
| A1 | Appendicitis | Unperforated | Acute appendicitis | Appendectomy | 32 | F | Unknown |
| A2 | Appendicitis | Unperforated | Acute appendicitis | Appendectomy | 32 | F | White |
| A3 | Appendicitis | Unperforated | Acute appendicitis | Appendectomy | 41 | M | Unknown |

### Follicular B cell regions show upregulation of B cell activation pathways and IgG genes, supporting a role for an antigen-driven adaptive immune response in appendicitis

In order to explore the changes taking place in the B cell follicle during appendicitis, gene expression was compared between appendicitis FB ROIs (n = 8) and normal FB ROIs (n = 12) (**Figure 1B**). Following differential expression analysis, 314 genes were significantly upregulated while 979 genes were downregulated in appendicitis (**Figure 1C**, 1D). Genes related to B cell lineage (*PAX5*), activation (*CD40, CD79A, CD22, LYN*), and isotype switching (*SWAP70, IGHM, IGHG2, IGHG4*) were among the top 30 most upregulated. All four IgG subclass genes had increased fold change in appendicitis, with *IGHG1* and *IGHG3* just shy of the p value cutoff, while *IGHG2* and *IGHG4* were significantly upregulated (**Figure 1E**). The gene encoding follicular dendritic cell secreted protein, *FDCSP*, was also upregulated in appendicitis B cell follicles (**Figure 1D**, **1E**). This protein is produced by FDCs and is known to bind preferentially to B cells activated by T-dependent anti-CD40 mechanism and was shown to regulate T-dependent antigen responses in the germinal center in a transgenic mouse model^17, 18^. Gene set enrichment analysis (GSEA) of FB segments also found enrichment in B cell activation pathways as well as antigen processing and presentation (**Figure 1F**, **Supplementary** Figure 1A). Taken together, these findings suggest there is increased B cell activation in the follicle during appendicitis, introducing the possibility for an antigen-driven adaptive immune response in this disease.

### Follicle-adjacent T cells show upregulation of T cell activation gene expression, supporting a role for the adaptive immune response in appendicitis

To perform an analogous exploration into T cell responses during appendicitis, gene expression was compared between appendicitis FT ROIs (n = 6) and normal FT ROIs (n = 3) (**Figure 1B**). Differential expression analysis revealed upregulation of 53 genes and downregulation of 12 in the FT regions (**Figure 1G**). Genes related to lymphocyte activation such as *CD69*, *JAK3, NAMPT, IL2RA* and *IFNGR1* were upregulated in appendicitis (**Figure 1H**). Genes related to complement activation (*CTSL)* and apoptosis regulation and oxidative stress (*CEBPD, CTSB, SOD2)* were also upregulated (**Figure 1H**). GSEA analysis revealed significant enrichment in T cell activation and differentiation pathways (**Figure 1I**). Additional pathways related to antigen presentation, innate immune activation, and immunoglobulin mediated immunity were also identified (**Supplementary** Figure 1B). This analysis supports the known role of innate immunity in appendicitis, while also providing evidence for T cell-mediated immunity.

### Single-cell RNA sequencing of appendiceal immune cells during acute appendicitis show a diversity of activated phenotypes in T and B cells

The immune mechanisms in appendicitis are not well understood, and little work has been done to characterize immune cell populations during disease. Based on the initial spatial transcriptomics analysis showing increased expression of genes related to adaptive immune activation, we wanted to further describe the T and B cell populations. To do this, we performed single cell RNA-sequencing (scRNA-seq) of immune cells isolated from the appendices of 5 donors undergoing appendectomy for appendicitis (**Table 2**). These cells were used for library preparation using the 5’ 10X Genomics workflow, which allows for sequencing of both total gene expression and V(D)J sequencing for immune repertoires (**Figure 2A**). After initial processing and quality control using the CellRanger pipeline, cell profiles were clustered and visualized using uniform manifold approximation and projection (UMAP), which resulted in 33 initial clusters. Clusters of less than 150 cells were excluded from all downstream analysis, as well as one cluster with high ribosomal gene content, resulting in 21 final UMAP clusters containing 47,179 cell profiles (**Figure 2B**). All clusters contained cells from each of the 5 donors. Major cell types identified included T cells, B cells, and one antigen presenting cell (APC) cluster containing macrophages. These three populations were relatively equal in proportion across all donors (**Figure 2C**). Of the T cell populations, the resting naïve/central memory (CM) CD4+ T cells (**C2**) was the most abundant, while tissue-resident memory (Trm) CD8+ T cells (**C13**) and CD161+ MAIT cells (**C16**) were the rarest relatively (**Figure 2D**). Within the B cells, the most abundant population was memory B cells (**C1**), while plasma cell clusters (**C8, C20**) were the rarest (**Figure 2E**).

**Figure 2.**
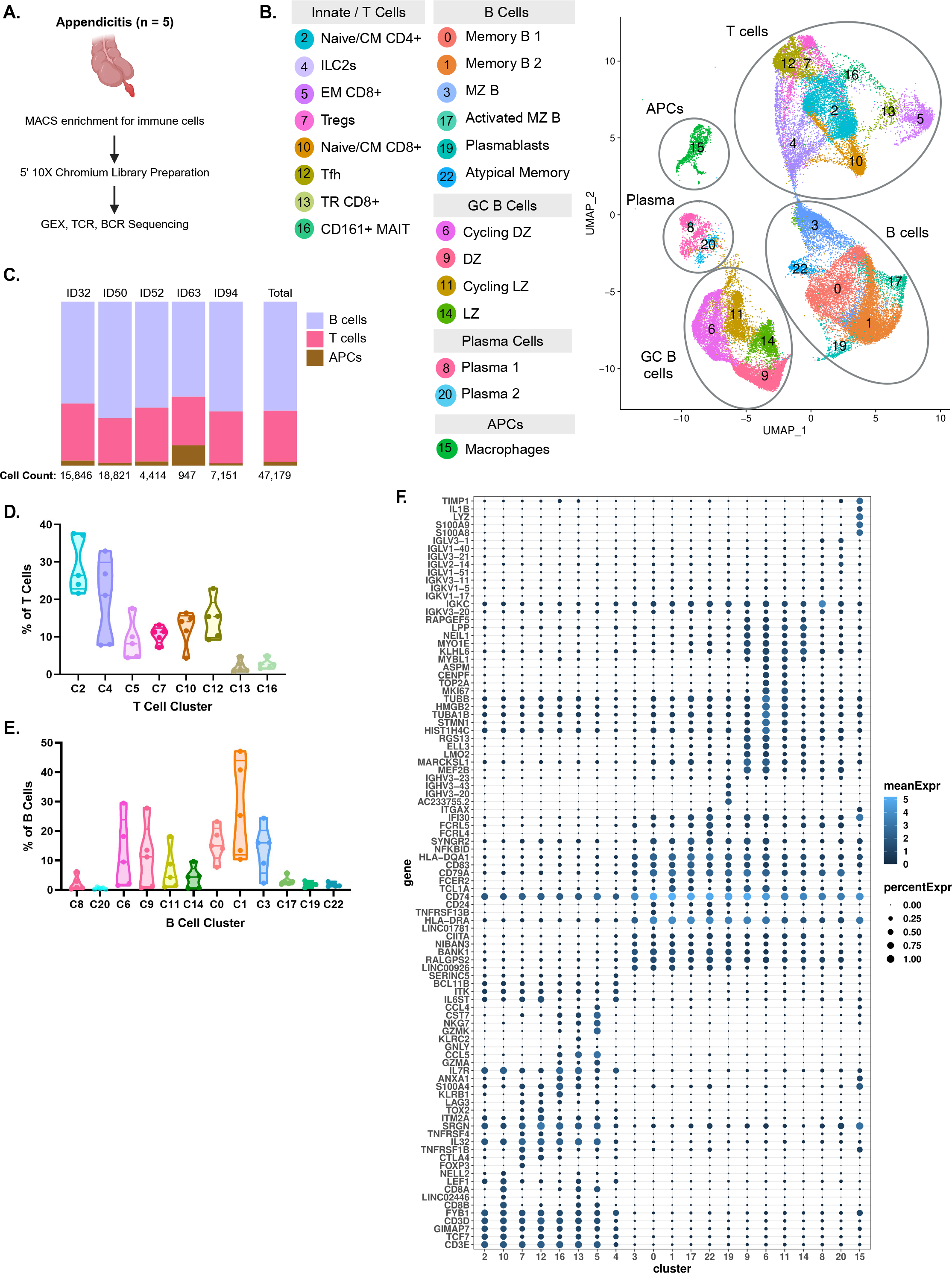
scRNA-seq analysis of appendiceal immune cells during appendicitis show diverse T and B cell sub-populations. **(A)** Workflow for 10X scRNA-seq sample collection and analysis from 5 appendicitis donors. **(B)** resulting UMAP of appendiceal immune cells after removal of clusters with high ribosomal gene content or containing fewer than 150 cells. Major cell groups are circled and labelled in the UMAP visualization, and all clusters annotations are listed. **(C)** Proportions of major cell types by donor. Total cells per donor are listed beneath the respective bar. **(D)** Truncated violin plot for frequencies of each T cell cluster as a percentage of all T cells, where each dot represents one donor. **(E)** Truncated violin plot for frequencies of each B cell cluster as a percentage of all B cells, where each dot represents one donor. **(F)** Bubble plot showing expression of key genes across clusters. Genes are shown the on the y axis, with clusters shown on the x axis. Bubble size corresponds to percent of cells within the cluster expressing that gene and are colored by mean gene expression. ***Abbreviations:*** UMAP = Uniform Manifold Approximation and Projection, GC = germinal center, APC = antigen presenting cell, CM = central memory, ILC = innate lymphoid cell, EM = effector memory, TR = tissue resident, MAIT = mucosal associated invariant T cell, MZ = marginal zone, DZ = dark zone, LZ = light zone

**Table 2.** Single-cell RNA sequencing cohort demographic and clinical characteristics.

| Donor ID | Study Group | Severity | Diagnosis/Indication | Surgery | Age | Sex | Race |
| --- | --- | --- | --- | --- | --- | --- | --- |
| ID50 | Appendicitis | Unperforated | Acute appendicitis | Appendectomy | 13 | F | Black |
| ID52 | Appendicitis | Unperforated | Acute appendicitis | Appendectomy | 15 | F | Unknown |
| ID63 | Appendicitis | Unperforated | Acute appendicitis | Appendectomy | 17 | M | Multiracial |
| ID32 | Appendicitis | Perforated | Acute appendicitis | Appendectomy | 8 | M | Black |
| ID94 | Appendicitis | Perforated | Interval appendicitis | Appendectomy | 7 | M | Unknown |

Eight of the clusters were grouped into the T cell portion of the UMAP. Of these, three clusters were the major CD8+ populations: resting naïve/CM CD8+ T cells (**C10**) (*TCF7, IL7R, CCR7, SELL*), cytotoxic effector CD8+ T cells (**C5**) (*CCL5, GZMK, KLRG1, PRF1*), and Trm CD8+ T cells (**C13**) (*ITGA1, ITGAE, RUNX3, CXCR6*). Another three clusters comprised the major CD4+ T cell clusters: resting naïve/CM CD4+ T cells (**C2**) (*TCF7, IL7R, LEF1, CCR7*), T regulatory cells (Tregs) (**C7**) (*FOXP3, CTLA4, IKZF2, IL2RA*), and T follicular helper cells (Tfh) (**C12**) (*PDCD1, ICOS, BCL6*) (**Figure 2B**, **2F, Supplementary** Figure 2A). Cluster 16 (**C16**) was a CD161+ (*KLRB1*) T cell cluster enriched with MAIT cell alpha chains. This cluster was tissue-resident (*ITGAE, CD69*) and expressed activation genes such as *NFKBIA, CD40LG,* and *TNF* (**Figure 2F**, **Supplementary** Figure 2A). Finally, cluster 4 (**C4**) while it had some TCR calling, was predominantly an innate lymphoid cell (ILC) cluster. Several transcription factors known to play a role in maintaining the ILC2 lineage were expressed (*TCF7, IKZF1, ETS1, RORA, ITK*) (**Figure 2F**, **Supplementary** Figure 2A). As ILC2s are known to play a role in tissue repair, it is likely they are recruited in appendicitis to respond to damage signals early in the inflammatory process.

Of the 12 B cell clusters, 2 were plasma cells, 4 were germinal center cells, and the remaining 6 were memory and innate-like populations (**Figure 2B**). Plasma clusters (**C8, C20**) were identified on the basis of *MZB1, XBP1, CCR10,* and *JCHAIN* expression. Germinal center clusters (**C6, C9, C11, C14**) expressed *BCL6* and *CD38*. Actively cycling clusters (**C6, C11**) expressed cell cycle genes such as *MKI67, PCNA, TOP2A*. Dark zone clusters (**C6, C9**) were differentiated from light zone (**C11, C14**) by *AICDA* expression. Additional B cell clusters included memory B cells (**C0, C1**) (*IGHM, IGHD, MS4A1, CD19, CD74*) and marginal zone B cells (**C3**) (*NOTCH2, CR1, CR2*).

Smaller clusters also included activated marginal zone B cells (C17), plasmablasts (C19) (*IGHV- genes, MZB1, IGHA1, IGHG1/3)*, and atypical memory cells (C22) (*FCRL4, FCRL5, ITGAX ENTPD1)* (**Figure 2F**, **Supplementary** Figure 2B).

The appendicitis scRNA-seq cohort overall identifies subsets of both activated T and B cells within the appendix. Within the T cell compartment the CD8+ effectors (**C5**) and CD161+ MAIT cells (**C16**) express genes related to cytotoxicity and activation, respectively. The B cell compartment contains relatively large clusters of germinal center cells. The germinal center is considered the major site of BCR maturation and development of high affinity antibody-producing plasma cells. Notably, germinal center reactions are also dependent on Tfh cells, which were identified in the CD4+ compartment.

### Repertoire analysis of appendicitis BCRs reveals expanded clonotypes in germinal center and plasma cell clusters with increased mutational frequency and IgG usage

In both the spatial transcriptomic analysis and scRNA-seq analysis, there was evidence for T and B cell activation. As adaptive immunity is generally dependent on antigen recognition via TCRs and BCRs, we performed repertoire analysis of both major cell types. In the B cell compartment, we hypothesized that there would be evidence of germinal-center dependent processes resulting in IgG+ B cells and plasma cells with increased mutational frequencies, based on our observation of increased IgG gene expression in appendicitis samples compared to normal appendices.

Amongst total BCRs, the majority of clones were IgM+, which accounted for approximately half of all BCR clones. The remaining clones were close to evenly split between IgA+ and IgG+, with very rare IgD+ clones (**Figure 3A**). Of the IgG+ clones, the majority were IgG1, which is reasonable as this is the most abundant IgG subclass. IgG2 and IgG3 made up approximately 20% and 10% of the remaining clones respectively, while IgG4+ clones were rare (**Figure 3B**). These IgG+ clusters predominantly mapped to germinal center clusters C6 and C9, along with the C8 plasma cell cluster (**Figure 3C**). Two donors had a majority of their IgG+ clones map to a memory B cell cluster (C0) (**Figure 3C**). Interestingly, these two donors had limited clonal expansion compared to the other three donors, highlighting (1) the heterogeneity in immune responses to appendicitis and (2) the possibility that these patients underwent appendectomy early enough in their disease course that they had not mounted a substantial humoral response. To get a global understanding of which BCR isotypes mapped to each cluster, we used a chord diagram to visualize movement of isotypes between clusters (**Figure 3D**). From this diagram it is obvious that memory cluster C1 is predominantly IgM+. The next two largest clusters, germinal center clusters C6 and C9 showed approximately half of BCRs were IgA+, one quarter IgM+, and the remaining quarter IgG+. The larger of the two plasma clusters, C8, was approximately split between IgA+ and IgG+ BCRs (**Figure 3D**). To quantify which clusters had enrichment of IgG+ BCRs, percentage of IgG of all BCRs within each cluster were plotted by donor and compared to the overall frequency of IgG. With this method we found IgG was significantly enriched in both plasma cell clusters (C8, C20) (**Figure 3E**). IgG was slightly increased in the germinal centers although this was not significant, and nearly absent in memory B clusters, suggesting that IgG+ BCRs first begin to appear in the germinal center, where mature cells eventually leave as they become plasma cells.

**Figure 3.**
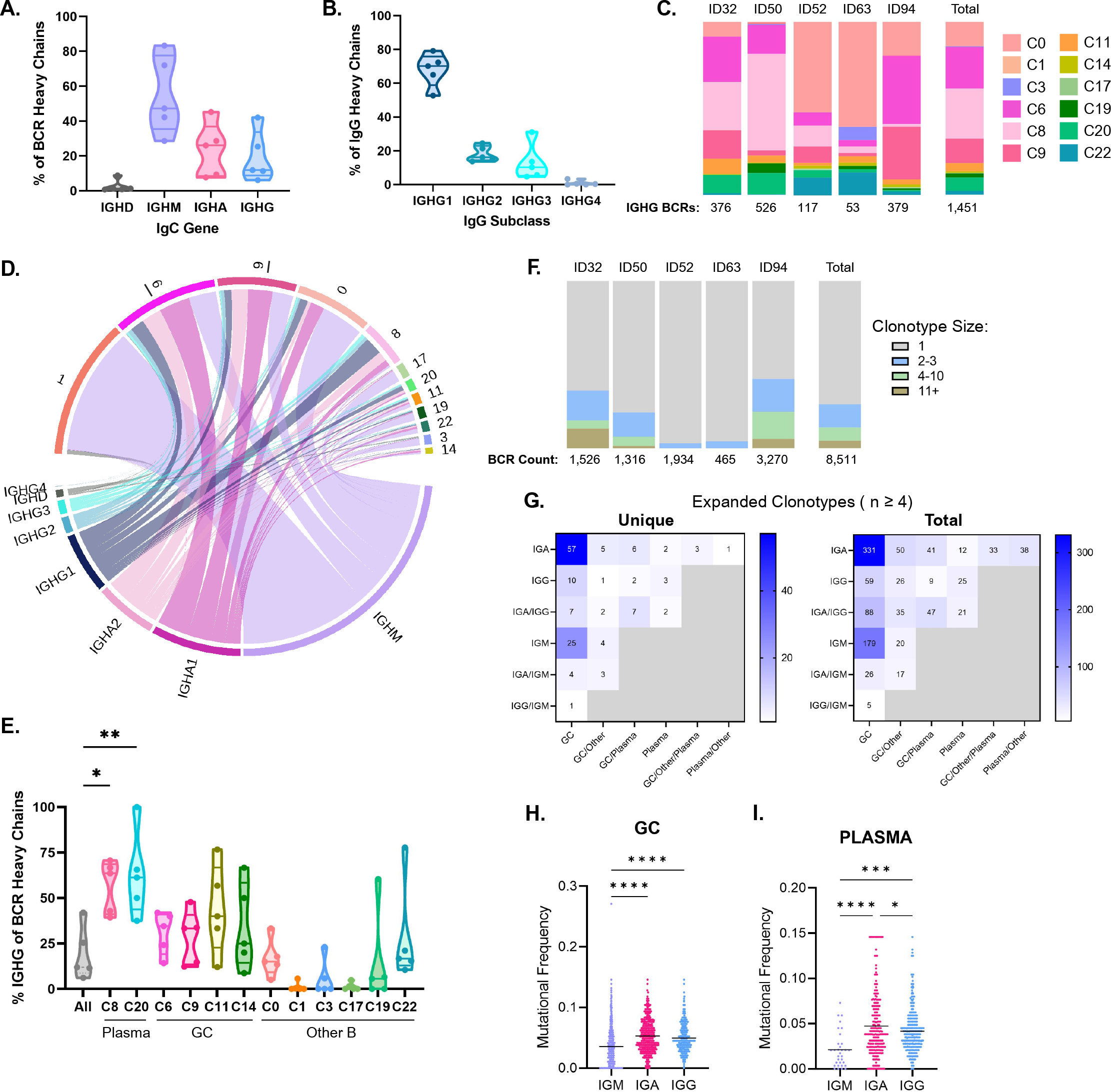
BCR repertoire analysis in appendicitis reveals expanded clonotypes in the germinal center and plasma cell clusters with increased mutational frequency and IgG usage. **(A)** Truncated violin plot of IgC gene usage as a percentage of BCR heavy chains, each dot represents an individual donor, median and quartiles are shown for each group. **(B)** Truncated violin plot of IgG subclass usage as a percentage of IgG heavy chains, each dot represents an individual donor, median and quartiles are shown for each group. **(C)** Bar plots showing relative frequencies of B cell clusters as a proportion of IgG+ heavy chains. Each bar plot represents one donor with IgG BCR counts listed below. **(D)** Chord diagram showing distribution of IgC genes across B cell clusters. Heavy chain constant regions are listed along the bottom, while cluster numbers are distributed across the top of the graph. Graph shows total BCR heavy chains from the appendicitis cohort. **(E)** Truncated violin plots showing the percentage of IgG+ BCRs across B cell clusters. Each dot represents one donor. Clusters were compared to overall IgG percentage (“All”) by ANOVA with multiple comparison testing. Significant comparisons are denoted by adjusted p-value ≤ 0.05 = *, ≤ 0.01 = **. **(F)** Bar plots showing relative proportions of total expanded BCR clonotypes across donors. Each bar represents one donor, with BCR heavy chain counts denoted below. **(G)** Heatmaps of BCR clonotypes with size ≥ 4, showing distribution of clones across BCR cluster group (x axis), and IgC gene (y axis). Clonotypes present in two or more groups are listed jointly. Left panel shows unique clonotype counts, right panel shows total BCR counts. **(H)** Mutational frequency plotted by C-gene usage in the germinal center compartment. Significance testing by ANOVA was performed with multiple comparison testing (****, p ≤ 0.001, ***, p ≤ 0.01, *, p ≤ 0.05). **(I)** Mutational frequency plotted by C-gene usage in the plasma cell compartment. Significance testing by ANOVA was performed with multiple comparison testing (****, p ≤ 0.001, ***, p ≤ 0.01, *, p ≤ 0.05).

As BCRs recognize their cognate antigen, they begin to proliferate. This process is complicated by the introduction of mutations via somatic hypermutation, however clones likely descended from the same lineage can be computationally predicted and annotated as being a single clonotype. We utilized the enclone annotation feature within the CellRanger pipeline to measure clonal expansion of the BCRs. All five donors had some clonal expansion observed (n > 2), with three of the donors having larger clone sizes (n ≥ 4) (**Figure 3F**). Next, expanded clones of size ≥ 4 were categorized by their isotype as well as UMAP location. Clonotypes with multiple isotypes were jointly labelled. UMAP locations were broken down by plasma cells (C8, C20), germinal center (C6, C9, C11, C14), and other (C0, C1, C3, C17, C19, C22) again with clonotypes in multiple groups jointly labelled. The resulting heatmap of clonotype mapping is shown in **Figure 3G**. From this map it is apparent that the largest number of unique clonotypes are present in the germinal center, with a smaller fraction of these expanding clones moving into the plasma compartment. IgA was the most common isotype for expanded clones, but clones jointly expressing IgA and IgG as well as IgG alone were also present in plasma cell clusters. When looking at mutational frequencies between Ig classes, both IgA and IgG had greater mutation rates than IgM in the germinal center and plasma cell compartments (**Figure 3H**, **3I**). These increased mutation rates indicate somatic hypermutation is taking place in the germinal center, likely leading to higher affinity IgA and IgG antibodies during appendicitis.

### Cytotoxic CD8+ T cells show evidence of TCR-independent bystander activation in appendicitis

To continue the investigation of the adaptive immune response in appendicitis, we next focused on the T cell receptor (TCR) repertoire. After initial quality control, TCRs were annotated as CD4+ or CD8+ on the basis of gene expression for their respective co-receptors or based on cluster assignment. Afterwards unpaired sequences were removed, resulting in 2,176 CD8+ TCRs (**Figure 4A**). The three major CD8+ T cell clusters were cytotoxic effectors in C5, resting naïve and memory cells in C10, and tissue-resident memory cells in C13. All 5 donors had TCRs present in the major CD8 clusters.

**Figure 4.**
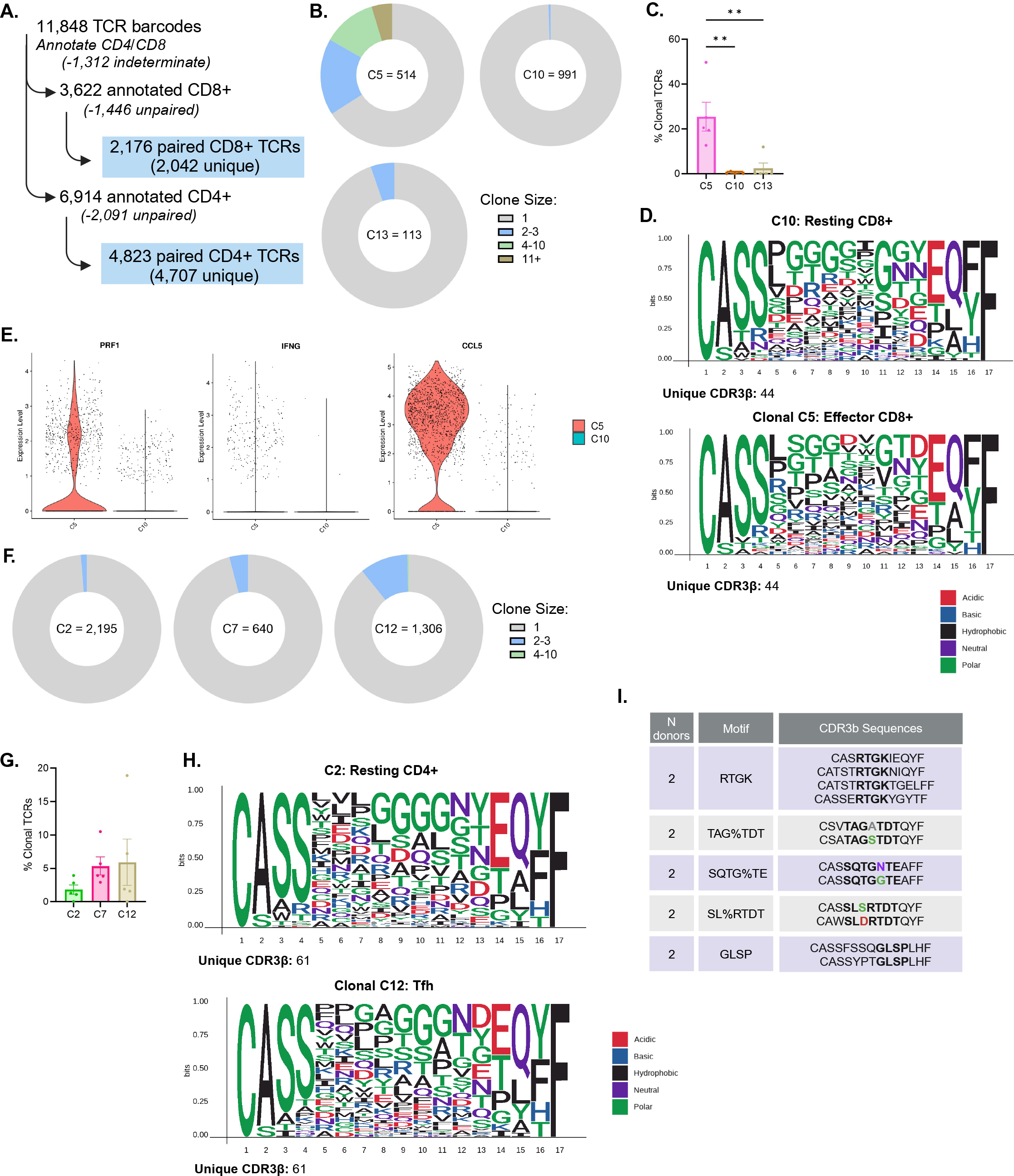
TCR repertoire analysis reveals likely bystander activation of conventional CD8+ T cells while CD4+ T-follicular helper cells show some evidence of antigenic drive. **(A)** Workflow to identify paired TCRs with CD4+ and CD8+ annotations for downstream analysis. **(B)** Pie charts showing clonal expansion of total TCRs for each of the three major CD8+ T cell clusters (C5, C10, and C13). Number of TCRs are included within each chart. **(C)** Bar graph showing percent of expanded clones across CD8+ T clusters, with each dot representing one donor. Comparisons between clusters were performed using ANOVA with multiple comparison testing, with significance noted as adjusted p value ≤ 0.01 = **. **(D)** Multiple sequence alignment of CDR3β sequences for length 12-17. Top panel shows relative amino acid use in 44 randomly selected sequences from the resting CD8 TCR subset (C10), bottom panel shows relative amino acid use in the 44 unique clonally expanded sequences from the effector CD8 subset (C5). **(E)** Violin plots showing gene expression by CD8+ T cell cluster for selected genes known to play a role in bystander activation. Significance was determined by ANOVA with multiple comparison testing. **(F)** Pie charts showing clonal expansion of total TCRs for each of the three major CD4+ T cell clusters (C2, C7, and C12). Number of TCRs are included at the center of each chart. **(G)** Bar graph showing percent of expanded clones in the CD4+ T cell clusters, with each dot representing one donor. Percent of clonal expansion was non-significant by ANOVA testing. **(H)** Alignment of 500 randomly selected CDR3β sequences from resting CD4+ TCRs (top) and T-follicular helper CD4+ TCRs (bottom) for sequences 12-16 amino acids in length. **(I)** Summary of GLIPH2.0 results using all CD4+ Tfh TCRs as input. Clusters with 2 or more donors were included in the summary, motifs in the CDR3β sequences are bolded, with distinct residues highlighted.

A key indicator of T cell antigen recognition and activation, we first assessed clonal expansion across CD8+ T cell clusters. As expected, clonal expansion in resting cells (C10) was minimal, while the effector subset (C5) had the greatest degree of expansion, both in terms of larger clone sizes and overall percentage of clonal expansion (**Figure 4B**, **4C**). Moving forward we focused on comparison of the effector TCRs (n = 514) versus resting TCRs (n = 991) to look for evidence of antigenic drive in the effector subset. However, we did not observe any significant differences in Vβ gene usage or CDR3β sequence length, nor was there enrichment for specific Vβ-Jβ or Vβ- Vα gene pairings in effector cells (**Supplementary** Figure 3A-D). To try and maximize any motif enrichment in the effector cluster, CDR3β sequences from expanded clones between 12-17 amino acids in length (n = 44) underwent multiple sequence alignment. We used a randomly selected subset of sequences of the same length from the resting CD8 population for comparison. Alignment and visualization of relative amino acid use did not show any obvious enrichment in the CDR3β motifs (**Figure 4D**). Residues between positions 5-13 showed minimal enrichment in specific amino acids, suggesting that TCRs in this subset were random, similar to what would be expected in the resting population. Next, we employed the GLIPH2.0 algorithm, which uses CDR3 sequences and Vβ gene usage to predict TCRs with likely similar reactivity^19^. However, inputting the effector CD8 TCRs resulted in no clusters with multiple donors, and single donor clusters consisted of a single shared CDR3β sequence, further indicating a lack of antigenic drive in this population.

The lack of TCR sequence enrichment in the CD8 compartment led us to examine the possibility of TCR-independent bystander activation. This mechanism has previously been described in multiple microbial infections in both mice and humans^20, 21^. We compared gene expression for known effectors in bystander activation across CD8+ T cell clusters. Expression of *PRF1, IFNG,* and *CCL5* were all significantly increased in the effector CD8+ population (C5) compared to the resting CD8+ population, supporting that these cells have the potential to contribute to inflammation during appendicitis (**Figure 4E**). During bystander activation, pro-inflammatory cytokines such as IL-15 and IL-18 are sufficient to activate memory CD8+ T cells, leading to perforin and granzyme production. In our single cell dataset, a small cluster of epithelial cells was positive for *IL15* expression, further supporting the possibility for bystander activation in appendicitis.

### TCRs from the Tfh CD4+ compartment show some evidence of antigen-driven expansion and activation in appendicitis

To investigate the possibility of antigen-driven activation in the CD4+ T cells, the same workflow was completed to annotate TCRs and remove unpaired sequences, resulting in a total of 4,823 CD4+ TCRs for downstream analysis (**Figure 4A**). There were three major CD4+ clusters, resting naïve/memory cells (C2), Tregs (C7), and Tfh (C12). Similar to the CD8 T cells, our first step was to investigate the extent of clonal expansion in the CD4+ TCR compartment. CD4 repertoires are generally less expanded than CD8s, which was in line with our observations from the appendicitis samples. The greatest degree of expansion was observed in the Tfh cluster (C12), accounting for 10.8% of total TCRs in this cluster (**Figure 4F**). However, percent of expanded clones did not reach significance (**Figure 4G**). Because the Tfh cluster had the greatest degree in clonal expansion, and because of their role in supporting B cell activation and maturation in the germinal center, we focused on comparison of the Tfh TCR compartment (n = 1,306) compared to resting TCRs (n = 2,195). Once again, there was no observed enrichment of Vβ gene usage, Vβ-Jβ pairing, Vβ-Vα pairing, or differences in CDR3β sequence length between the two clusters (**Supplementary** Figure 4A-D). We repeated the motif sequence alignment analysis for the CD4 TCRs. Again, to try and enrich for antigen-specific TCRs potentially relevant in disease, we selected the clonally expanded CDR3β sequences from the Tfh cluster (n = 61) with sequence length between 12-17 amino acids. A random selection of sequences from the resting CD4+ TCRs was used for comparison (n = 61). Within the clonally expanded Tfh TCRs there was still substantial diversity in relative amino acid use, with some variation in positions 5-8 compared to the resting cohort, along with enrichment of aspartic acid at position 13 (**Figure 4H**). All Tfh TCRs were also input into GLIPH, this time with multiple clusters shared between donors. These included both local and global sequence motifs, suggesting the possibility of shared epitope specificity within this cluster (**Figure 4I**). These results and the presence of greater clonal expansion in the Tfh subset supports the presence of antigen-driven activation of these cells, facilitating the maturation of B cells in the germinal center.

**Appendicitis is characterized by outgrowth of opportunistic pathogens such as Parvimonas that are positively correlated with oral biofilm-forming genera such as Fusobacterium.**

After defining the immune cell populations and adaptive repertoires in the appendix during appendicitis, it led us to consider what environmental factors which may also contribute to appendicitis pathology. Microbial infections have been implicated in appendicitis in the literature, and the appendix contains a dense microbiome, leading us to hypothesize that alterations in the microbiome are present during appendicitis and contribute to the inflammatory response during disease. To characterize changes in the appendicitis microbiome, we collected appendiceal swabs for 16S sequencing from 10 controls undergoing incidental appendectomy, and 12 appendicitis donors, which were further separated into perforated (n = 7) and unperforated (n = 5) subgroups. Demographic and clinical characteristics for this cohort can be found in **Table 3**. When comparing relative frequencies from the most abundant genera in the dataset, it was readily apparent that many of the appendicitis samples, in particular the perforated appendicitis cases, had an outgrowth of *Parvimonas* that was nearly absent from all control samples (**Figure 5A**). In line with prior literature, there was no significant difference in alpha diversity between the three study groups as measured by the Shannon diversity index (**Figure 5B**). However, beta diversity using the Bray-Curtis index was significantly different between study groups, and pairwise comparison found a significant alteration in community composition of perforated appendicitis samples as compared to controls (**Figure 5C**). These results indicate that while the appendicitis microbiome is not less diverse than those of control samples, the community composition is altered, especially in the perforated cases.

**Figure 5.**
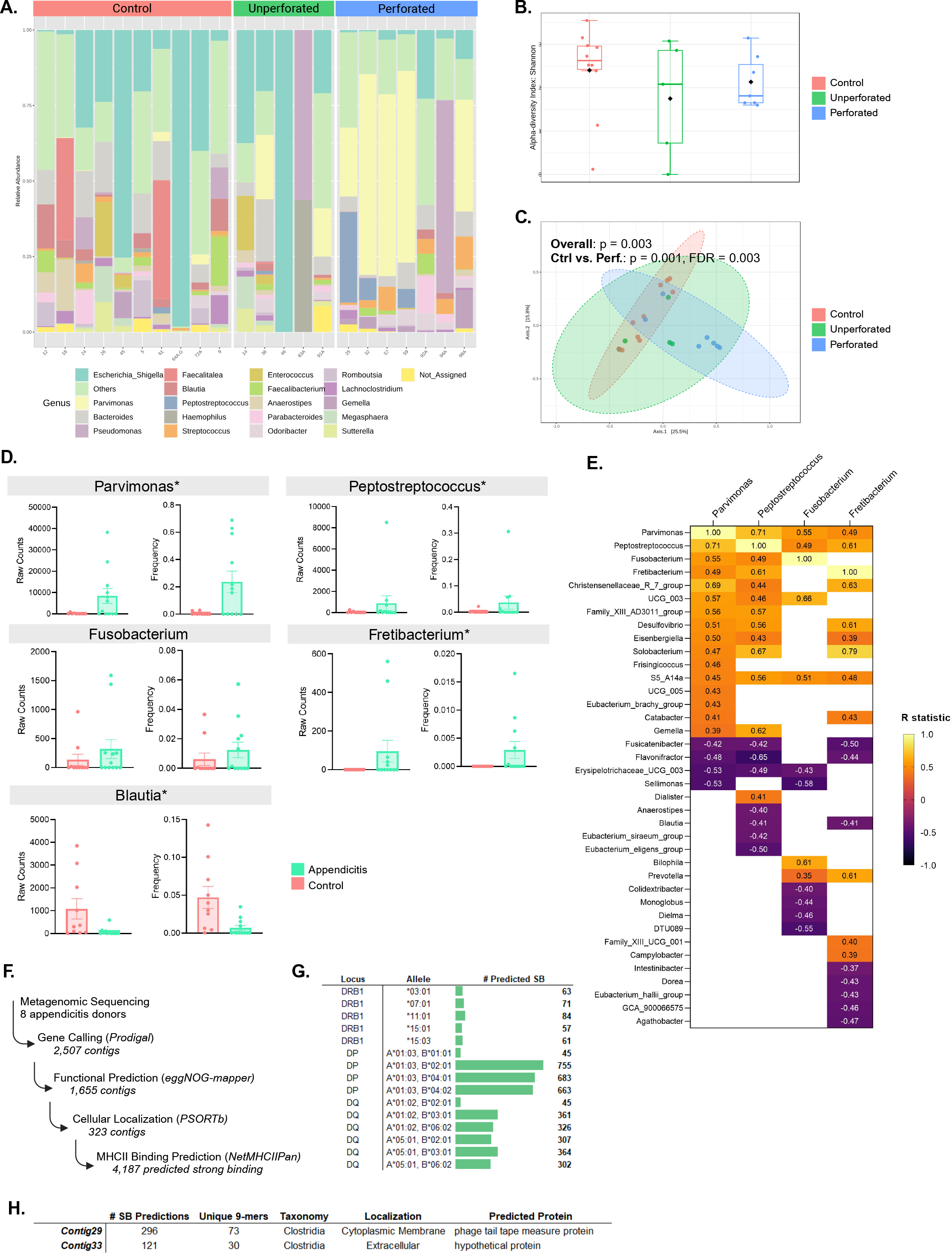
The appendiceal microbiome during perforated appendicitis is characterized by outgrowth of opportunistic pathogens which correlate with presence of oral biofilm-forming genera. **(A)** Relative abundances for the top 20 most abundant genera by donor are shown, grouped by controls and perforation status of appendicitis samples. **(B)** Alpha diversity of samples as measured by the Shannon diversity index; no significant difference was determined by ANOVA test. **(C)** Beta diversity was calculated using the Bray-Curtis index and visualized with PCoA plot. Significance testing was performed using PERMANOVA test along with pair-wise comparison using the Benjamin-Hochberg adjustment, with FDR cutoff of ≤0.05. **(D)** Raw counts and frequencies are plotted for selected genera. Labels with * were significant by edgeR analysis comparing controls versus appendicitis samples, with significance determined as an FDR cutoff of ≤0.05. **(E)** Correlation network analysis was performed using sparCC, with correlation coefficient cutoff ≥0.3 and p-value ≤0.05 for inclusion in the analysis. A heatmap of resulting correlations for select genera is shown. **(F)** Workflow for metagenomic sequencing analysis to identify potential microbial derived peptides in appendicitis. Appendicitis samples with DNA extract remaining following 16S analysis were re-sequenced, with subsequent gene calling and functional prediction (SB = strong binder as predicted by top 1% rank using NetMHCIIPan). **(G)** Table summarizing number of predicted strong binding sequences by MHC allele. Alleles were chosen as being the most frequent amongst representative U.S. Caucasian and African American populations. **(H)** Summary of top 2 most frequent contigs that were present in the strong binder results, along with number of unique 9-mer binding cores, cellular localization, and predicted protein product.

**Table 3.** 16S microbiome cohort demographic and clinical characteristics.

| Donor ID | Study Group | Severity | Diagnosis/Indication | Surgery | Age | Sex | Race |
| --- | --- | --- | --- | --- | --- | --- | --- |
| ID12 | Control | NA | Colonic adenocarcinoma | Cytoreduction | 61 | F | White |
| ID18 | Control | NA | Colonic adenocarcinoma | Right hemicolectomy | 80 | F | White |
| ID24 | Control | NA | Cecal adenocarcinoma | Right hemicolectomy | 56 | M | Black |
| ID26 | Control | NA | Colonic adenocarcinoma | Right hemicolectomy | 72 | F | White |
| ID45 | Control | NA | Colonic adenocarcinoma | Right hemicolectomy | 78 | F | White |
| ID5 | Control | NA | Neuroendocrine tumor | Right hemicolectomy | 56 | M | White |
| ID61 | Control | NA | Colonic adenocarcinoma | Right hemicolectomy | 78 | F | Unknown |
| ID64A .G | Control | NA | Tubular adenomas | Right hemicolectomy | 68 | M | Black |
| ID72A | Control | NA | Colonic adenocarcinoma | Total colectomy | 37 | M | Unknown |
| ID9 | Control | NA | Colonic adenocarcinoma | Right hemicolectomy | 65 | M | White |
| ID14 | Appendicitis | Unperforated | Acute appendicitis | Appendectomy | 13 | F | White |
| ID36 | Appendicitis | Unperforated | Acute appendicitis | Appendectomy | 32 | M | White |
| ID46 | Appendicitis | Unperforated | Interval appendicitis | Appendectomy | 13 | F | White |
| ID83A | Appendicitis | Unperforated | Acute appendicitis | Appendectomy | 23 | M | White |
| ID91A | Appendicitis | Unperforated | Acute appendicitis | Appendectomy | 11 | M | White |
| ID25 | Appendicitis | Perforated | Acute appendicitis | Appendectomy | 2 | M | White |
| ID32 | Appendicitis | Perforated | Acute appendicitis | Appendectomy | 8 | M | Black |
| ID57 | Appendicitis | Perforated | Acute appendicitis | Appendectomy | 5 | F | White |
| ID59 | Appendicitis | Perforated | Acute appendicitis | Appendectomy | 16 | F | White |
| ID92A | Appendicitis | Perforated | Acute appendicitis | Appendectomy | 15 | F | White |
| ID94A | Appendicitis | Perforated | Interval appendicitis | Appendectomy | 7 | M | Unknown |
| ID98A | Appendicitis | Perforated | Acute appendicitis | Appendectomy | 13 | F | White |

In order to identify which species were altered in appendicitis, we performed edgeR differential expression analysis. Comparison of the appendicitis and control group yielded 22 significant results using a false-discovery rate cutoff ≤ 0.05. Within the significant genera, we found opportunistic pathogens such as *Parvimonas* and *Peptostreptococcus* were significantly increased in appendicitis (**Figure 5D**). A previous publication also reported increased *Parvimonas* specific to complicated appendicitis that was associated with oral bacterial pathogens such as *Fusobacterium*^10^. While increased *Fusobacterium* has repeatedly been associated with appendicitis in the literature, we observed a trend towards an increase in this genus, although it did not reach significance^7, 22, 23^. However, another subgingival plaque genus, *Fretibacterium*, was significantly increased in appendicitis (**Figure 5D**).

We next wanted to understand if any of the differentially expressed groups were associated with one another, indicating a “signature” of microbial dysbiosis in appendicitis. We performed SparCC network correlation analysis comparing appendicitis versus control samples, using cutoffs of ≥ ±0.3 correlation coefficient and p value ≤ 0.05 to be included. Within the resulting network, *Parvimonas, Peptostreptococcus, Fusobacterium,* and *Fretibacterium* were all positively correlated with one another (**Figure 5E**). This subset of microbes has all been isolated from subgingival plaque and are associated with periodontal disease. While a specific mechanism is yet to be identified, these genera are known to be present in multiple disease states, providing a rationale for further studies looking at the role for invasive biofilms in appendicitis.

### Identification of MHC-II restricted immunogenic peptides from the appendiceal microbiome during appendicitis

The microbiome likely contributes to appendicitis pathogenesis, as evidenced by efficacy of antibiotics in treating some cases. Our study and previous literature have identified changes in microbial communities during disease, leading us to wonder whether there are microbial antigens relevant to appendicitis. While the majority of changes we observed in the adaptive immune response were in the B cell compartment, prediction of BCR-epitope binding remains extremely challenging. However, our analysis revealed germinal center activation, a process which is not only antigen-dependent but also T cell dependent. In the germinal center, Tfh cells aid in B cell activation via recognition of antigen in an MHC-II restricted manner. Our analysis also identified some potentially shared reactivity in TCR sequences during GLIPH2.0 analysis, as well as some clonal expansion, suggesting antigen-dependent activation in this compartment.

Computational prediction of MHC-II restricted microbial-derived epitopes has been described in the literature, demonstrating feasibility of this technique^24^. To try and identify predicted CD4+ T cell epitopes from the appendicitis microbiome, we submitted a subset of appendiceal swabs for metagenomic sequencing. This cohort consisted of 3 unperforated and 5 perforated samples, with additional characteristics available in **Table 4**. We employed a similar workflow as previously described to identify potential microbial-derived epitopes using the metagenomic data from the nine appendicitis donors (**Figure 5F**). By performing gene calling, functional annotation of the likely gene product, and prediction of cellular localization, we were able to identify products more likely to be antigenic. These sequences are more likely to be derived from extracellular or membrane proteins. This resulted in identification of 323 contigs for input into NetMHCIIPan, along with the most frequent MHC alleles at each locus based on representative U.S. populations. Using a 1% rank cutoff, 4,187 predicted strong binding 15 amino acid-length peptides were identified from these contigs. The majority of these strong-binders (SB) were predicted to bind to alleles at the DP locus, with relatively few results for the DRB1 locus (**Figure 5G**). To try and understand whether certain contigs had an outsized impact, we calculated how frequently each contig appeared in the SB dataset. Two sequences, Contig29 and Contig33 were by far the most abundant in the dataset (**Figure 5H**). When looking at the potential binding sequences, 73 and 30 unique 9-mer binding cores were identified in each contig respectively. Both were predicted to be derived from the *Clostridia* class. While Contig33 was a hypothetical extracellular protein, interestingly, Contig29 was predicted to be a tape measure protein from a *Clostridia*-derived bacteriophage (**Figure 5H**). This analysis demonstrates that there is high potential for microbial-derived antigens to contribute to antigen-driven immune responses, as thousands of peptides were identified from a relatively limited number of contigs and provides rationale for a deeper dive into identifying highly immunogenic bacterial epitopes in appendicitis.

**Table 4.** Metagenomic analysis microbiome cohort demographic and clinical characteristics.

| Donor ID | Study Group | Severity | Diagnosis/Indication | Surgery | Age | Sex | Race |
| --- | --- | --- | --- | --- | --- | --- | --- |
| ID36 | Appendicitis | Unperforated | Acute appendicitis | Appendectomy | 32 | M | White |
| ID46 | Appendicitis | Unperforated | Interval appendicitis | Appendectomy | 13 | F | White |
| ID63 | Appendicitis | Unperforated | Acute appendicitis | Appendectomy | 17 | M | Multiracial |
| ID25 | Appendicitis | Perforated | Acute appendicitis | Appendectomy | 2 | M | White |
| ID32 | Appendicitis | Perforated | Acute appendicitis | Appendectomy | 8 | M | Black |
| ID57 | Appendicitis | Perforated | Acute appendicitis | Appendectomy | 5 | F | White |
| ID59 | Appendicitis | Perforated | Acute appendicitis | Appendectomy | 16 | F | White |
| ID60 | Appendicitis | Perforated | Interval appendicitis | Appendectomy | 11 | F | White |

### Evidence of epithelial barrier integrity disruption in appendicitis identifies a likely route for microbial tissue invasion

Microbial entry into the tissue at mucosal sites is usually tightly regulated by mucus production, the epithelial barrier, and the host immune system. However, when epithelial barrier integrity is compromised, microbes can invade the tissue, leading to damage and inflammation. *F. nucleatum* was previously implicated in tissue invasion in appendicitis, and the results of our microbiome analysis found dysbiosis in appendicitis and rise of opportunistic pathogens^25^. To investigate whether there was evidence of epithelial barrier integrity disruption in appendicitis, we analyzed spatial transcriptomic data from epithelial cell segments in the appendix. Representative ROIs are shown from both an appendicitis sample and normal control, outlined and labelled as “EPI” (**Figure 6A**). Nine ROIs were collected from three appendicitis donor tissues, and 12 ROIs were collected from 4 normal donor tissues (**Figure 6B**). Differential expression analysis revealed 54 upregulated genes and 9 downregulated genes out of 8,805 total genes detected (**Figure 6C**, **6D**).

**Figure 6.**
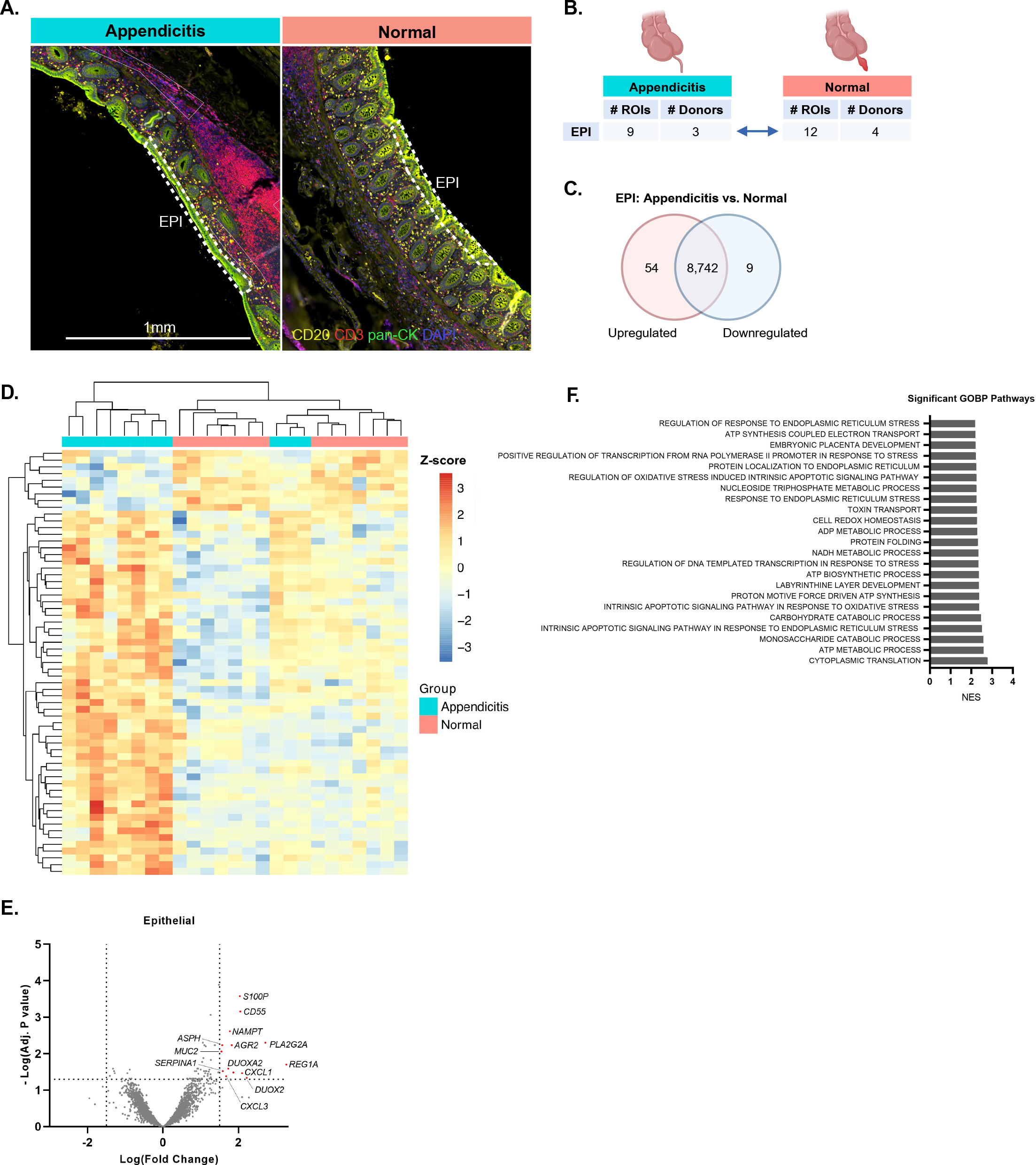
Appendiceal epithelial cells upregulate transcriptional pathways related to cell stress and loss of barrier integrity, highlighting the role of barrier disruption in appendicitis pathology. **(A)** Representative ROIs for epithelial cells are outlined in white dashed lines and labelled “EPI” from both appendicitis and normal FFPE tissue sections. **(B)** Workflow for spatial transcriptomic analysis, showing the summary of ROIs collected and number of donors for EPI segments. **(C)** Summary of significant results from DE gene analysis, based on adjusted p-value cutoff of ≤0.05. **(D)** Heat map showing all significant DE genes, colored by Z score. **(E)** Volcano plot of DE genes, based on adjusted p value cutoff of ≤0.05 and log(FC) ≥ ±1.5. Upregulated genes are shown in red. **(F)** Significant GSEA pathways as determined using an FWER p-value cutoff of ≤0.05. Pathways are ordered by normalized enrichment score (NES).

Both *DUOX2* and *DUOXA2* were significantly increased in the epithelium of appendicitis tissue (**Figure 6E**). *DUOX2,* along with its regulatory counterpart *DUOXA2*, encodes a protein which produces hydrogen peroxide in a microbiota-dependent fashion^26^. Mice with increased activation of DUOX2 showed increased bacterial translocation which led to increased pro-inflammatory cytokine signaling^27^. Increased expression of this gene has been found in patients with inflammatory bowel disease, and conditional knockout of *Duox2* in intestinal epithelium protected mice against DSS-induced colitis^28^. *REG1A* was also increased in the epithelium of appendicitis patients in our dataset (**Figure 6E**). This gene is also upregulated in inflammatory bowel disease, where it is expressed in response to IL-6/IL-22 mediated JAK/STAT3 signaling and acts as a protective factor to inhibit inflammatory responses^29^. Results of GSEA analysis on the epithelial segments also found multiple pathways related to cell stress and apoptosis (**Figure 6F**). These changes in gene expression in the epithelium of appendicitis samples supports a loss of epithelial barrier integrity and the potential for microbes to invade the nearby tissue and contribute to disease pathology.

## Discussion

Our study provides a high-resolution characterization of the appendix during appendicitis, one of the most common abdominal emergencies. Our spatial transcriptomic analysis of the lymphoid follicles found increased IgG genes and B cell activation, indicating an antigen-driven response, as IgG is canonically thought to be produced in a T cell- and antigen-dependent fashion. scRNA-seq and BCR repertoire analysis identified enrichment of IgG in the germinal center and plasma cell compartment, along with increased somatic hypermutation in this compartment, which suggests an antigen-driven process is taking place. We also found evidence of increased T cell activation adjacent to B cell follicles. Single-cell RNA-sequencing confirmed the presence of activated CD8+ T cells, which we propose are driven by bystander activation, as there was no clear evidence of antigenic drive based on TCR repertoire analysis. However, there was some evidence of antigenic drive in the T-follicular helper CD4+ cells by GLIPH2.0 analysis. This led us to pursue potential sources of antigen in the appendix. The most intuitive source is the microbiome, which is the densest site of biofilm in the GI tract. 16S analysis identified marked dysbiosis in the appendix during appendicitis, and spatial transcriptomics also found disruption of epithelial barrier integrity, providing rationale for invasion of microbes into the tissue during appendicitis. This led us to perform exploratory analysis of potential microbial derived-MCH II restricted peptides using a subset of metagenomic sequences from appendicitis samples. As BCR epitopes remain technically difficult to predict, this gives us a hint as to which peptide sequences the CD4+ T-follicular helper cells may be recognizing to assist in germinal center reactions.

A limitation of this study was difficulty in obtaining healthy control appendices. Our single-cell cohort included only young donors with appendicitis, as it was not feasible to obtain healthy appendix tissue from similarly aged donors. In our 16S microbiome cohort, the controls were generally patients undergoing surgery for colorectal cancer where the appendix was uninvolved in disease and were much older than the appendicitis cases. As a result, age may be a confounding factor in the microbiome analysis. However, we identified similar features in our cohort as to what has previously been reported in the literature, which supports the validity of our results despite the differences in age.

A common finding in appendicitis is the presence of subgingival plaque genera that are associated with periodontal disease. In line with this observation, we also found increased presence of *Fusobacterium* and *Fretibacterium*. Subgingival plaque is a biofilm, and the appendix has the densest biofilm formation in the gastrointestinal tract. Our network correlation analysis found that *Fusobacterium, Fretibacterium, Parvimonas,* and *Peptostreptococcus* were all positively correlated with one another in appendicitis. Interestingly, *Parvimonas micra* and *Fusobacterium nucleatum* have been shown to have synergistic biofilm formation, and *Peptostreptococcus micros* has been shown to co-aggregate with *F. nucleatum*^30, 31^. Additional species assignment using the Silva taxonomy database for our 16S data found that 69% of *Fusobacterium* sequences belonged to *F. nucleatum,* with the remainder unassigned. *Parvimonas micra* and *Peptostreptococcus stomatis* were the predominant species in their respective genera. These three species, *F. nucleatum, P. micra,* and *P. stomatis* have previously been associated with colorectal cancer (CRC), and were proposed as part of a biomarker panel for identifying CRC patients^32, 33^. While the specific mechanism remains undefined, these species have been implicated in multiple inflammatory conditions, including appendicitis, periodontitis, and cancer, and provides a rationale for further studies looking at the role of biofilm formation during disease.

It is possible that these genera form appendiceal biofilms that are better able to survive in the inflammatory conditions of appendicitis, leading to the relative outgrowth we observed in the 16S cohort. Periodontal disease has previously been described as having inflammophilic bacterial communities, meaning the species present thrive under inflammatory conditions, allowing them to outcompete other species^34, 35^. *Parvimonas* and *Fusobacterium*, as part of microbial cohorts, were shown to reduce neutrophil killing via dampening of reactive oxygen species production, and produced proteases to degrade immunoglobulins and complement proteins, reducing phagocytosis by host immune cells^35^. Both *Parvimonas* and *Fusobacterium* have been shown to induce inflammatory cytokines like IL-6 and IL-8^35, 36^. *Fusobacterium* was shown to invade the epithelial barrier in appendicitis^25^. In a model of periodontitis, this invasion was shown to be mediated via adhesin FadA, and found negative effects of bacterial invasion could be attenuated via treatment with piperlongumine or fisetin^36^. These studies highlight the importance of understanding how microbial dysbiosis can contribute to inflammation during appendicitis and may justify further studies to repurpose existing compounds for use in appendicitis.

Accurate grading of appendicitis severity was also a challenge for this study. We classified appendicitis cases as perforated versus unperforated upon gross examination, as a proxy for identification of complicated versus uncomplicated appendicitis. Complicated appendicitis is typically defined as gangrenous or perforated tissue, although some micro-perforations may require histological analysis and are not grossly apparent. Uncomplicated appendicitis cases show evidence of inflammation but no evidence of perforation. Differentiation between complicated and uncomplicated disease is imperfect, even with imaging, as symptoms of appendicitis are nonspecific and there are no useful biomarkers. This classification is clinically relevant, as it has been shown that some uncomplicated appendicitis cases can be treated by antibiotics alone. A recent review found that patients with imaging findings suggestive of complicated disease course, including appendiceal diameter ≥13 mm, presence of fecalith, or mass effect on computed tomography scans indicate increased risk of antibiotic failure^37^. However, identifying patients who can be successfully treated with antibiotics is an imperfect science, and up to 40% of treated patients will have recurrent appendicitis and require appendectomy in the future^13, 14^. Persisting challenges in identifying patients where antibiotic therapy is appropriate provides a rationale for further research into the disease mechanisms of appendicitis.

It is up for debate whether appendicitis is a progressive disease, where left untreated all cases will become complicated, or if these classifications represent two distinct diseases with distinct inflammatory mechanisms. In support of appendicitis as a progressive disease, retrospective studies have found that delaying surgery by greater than 24 hours increases the risk of perforation and complications^38^. Additionally, while overall incidence of appendicitis was decreased, perforation rates increased during the COVID-19 pandemic, presumptively due to treatment delays while patients and providers were trying to limit contact^39^. While the debate of appendicitis as a progressive disease versus distinct mechanisms is still an open question, we found evidence of increased B cell activation and IgG production in both perforated and unperforated samples in our transcriptomic data, supporting the role of the humoral response in both “types” of appendicitis.

The appendix has largely been neglected in scientific literature due to the assumption that it is a vestigial organ. While it is true the appendix is not necessary for life, that does not mean it serves no purpose. The appendix houses both dense lymphoid tissue and microbiota. Appendiceal lymphocytes and the microbiome are intimately linked. Follicle development coincides with bacterial colonization of the appendix in early human life^40^. The appendix is also a major producer of IgA, which helps to maintain microbial homeostasis at mucosal interfaces^3, 40–42^. Due to its narrow lumen and relative isolation from the fecal stream, the appendix has been hypothesized to act as a microbial reservoir that can re-seed the gut following infection or other intervention. Interestingly, bariatric surgery patients who underwent prophylactic appendectomy experienced reduced fecal microbial diversity compared to bariatric surgery patients whose appendices were left intact, supporting a role for the appendix in restoring the gut microbiome in times of microbial upset^43^. Additionally, inflammatory responses are not only of interest during appendicitis. The appendix has been associated with multiple diseases, including colorectal cancer, ulcerative colitis, and Parkinson’s disease, although via mechanisms unknown^44–48^. Additional research regarding the environment of the human appendix may reveal new insights into both the function of this organ as well as insights into the pathophysiology of multiple human diseases.

## Methods

### Spatial transcriptomic data collection and analysis

Tissue blocks were identified for appendicitis patients and age-matched controls who had undergone incidental appendectomy. Fresh FFPE sections were prepared by the Human Tissue Resource Center at UChicago and stored in the fridge prior to staining. Tissues were stained with anti-CD20 (clone ID: IGEL/773, 615nm channel), anti-CD3 (clone ID: C3e/1308, 666nm channel), and anti-pan-cytokeratin (clone ID: AE1/AE3, 532nm channel) antibodies in addition to DNA dye SYTO 13 (525nm channel) to identify regions of interest. ROIs were profiled for spatially indexed transcriptomics using the Human Whole Transcriptome Atlas following manufacturer instructions. Sequencing was performed by the Functional Genomics Core at UChicago using the NovaSeq6000 platform.

Raw count data was processed using the GeoMx Tools R package (v3.2.0). Segments with fewer than 1000 raw reads and sequencing saturation less than 50% were filtered out from the analysis. A probe was removed if (geometric mean of probe count from all segments / geometric mean of all probe counts from all segments) < 0.1 or if the probe was an outlier according to Grubb’s test in at least 20% of the segments. Segments with less than 5% of genes detected were removed, and genes detected in fewer than 10% of segments were removed. Counts were then normalized by the Q3 normalization.

Differential expression for each subset (“FB”, “FT”, “EPI”) was modelled using linear models with experimental factors as the predictor using limma^49^. The main factor of disease group (i.e. “Appendicitis” versus “Normal”) was used as a covariate. The *topTable* function was applied to calculate test statistics including the log fold-change and adjusted p-value for all genes. An adjusted p-value cutoff of ≤ 0.05 was used as a significance threshold for differential expression.

Gene set enrichment analysis (GSEA) was performed using total gene lists from each compartment using the PreRanked function within the GSEA software using default settings using the C5 Gene Ontology Biological Processes gene sets^50, 51^. T test values were used for rankings. GSEA pathway significance was determined using an FWER p-value cutoff of ≤ 0.05.

### Generating single cell suspension from appendiceal tissue

Appendicitis cases undergoing appendectomy were identified daily, with patient consent obtained by the Human Tissue Resource Center at UChicago. Appendiceal tissue sections were collected by Anatomic Pathology at UChicago and transported to the lab on ice in PBS (supplier, cat #). Appendix tissue was stored in MACS Tissue Storage Solution (Miltenyi Biotec cat. # 130-100- 008) until processing.

Tissue pieces were weighed and placed in a petri dish containing RPMI-1640 (Cytiva, #SH30027.1). Appendiceal sections were cut open along the orifice to reveal the lumen, and the outer serosal layer was peeled away using forceps and discarded. The mucosal layer was gently peeled or scraped into the media using forceps. The remaining submucosal tissue was placed in a conical tube containing fresh RPMI-1640 supplemented with digestion enzymes from the human Tumor Dissociation Kit (Miltenyi Biotec, #130-095-929) and chopped into small pieces using dissection scissors. Tissue was incubated at 37°C for 30 minutes with rotation to break down appendix tissue. Following digestion, both mucosal and submucosal cells were double filtered through 100µm and 70µm filters respectively and washed with fresh media for cell counting.

Afterwards, cells were washed once more in Milli-Q water containing 1X BSA (Miltenyi Biotec #130-091-376) and 1X Annexin V Binding Buffer (BD Biosciences, #556454) for magnetic enrichment. Cells were incubated in the dark on ice for 20 minutes with APC anti-CD235a (Biolegend, #349114), APC anti-EPCAM (Biolegend, #342208), and APC Annexin V (Invitrogen, #BMS306APC-100), washed, then followed by another incubation with anti-APC Microbeads (Miltenyi Biotec, #130-090-855). Negative enrichment to remove red blood cells, epithelial cells, and dying cells was performed using LS Columns (Miltenyi Biotec, #130-042-401) and the QuadroMACS Separator (Miltenyi Biotec, #130-090-976).

### Preparation of 10X single cell libraries

Following immune cell enrichment, cells were washed once more with 0.05% UltraPure BSA (ThermoFisher Scientific, #AM2616) and counted once more in preparation for the single-cell workflow. Cells were resuspended to the recommended cell density in the 10X 5’ v2 protocol. GEM generation was performed according to the manufacturer’s recommendations using the 10X Chromium controller. Single cell libraries were generated using the 5’ v2 reagent kits and associated protocols. Library QC was performed using the BioAnalyzer High Sensitivity DNA Kit (Agilent, #5067-4626) and run on the 2100 BioAnalyzer Instrument. Samples were pooled and submitted for sequencing at the UChicago Functional Genomics Core. Samples were sequenced with either the Illumina NovaSEQ-6000 or NovaSeqX.

### Single cell library QC and GEX analysis

Initial quality control of 10X single-cell RNA seq data was performed at the CRI Bioinformatics Core at UChicago using existing pipelines. Uniform manifold approximation and projection (UMAP) visualization was performed to visualize clusters. Clusters of less than 150 cells or with top genes consisting primarily of rRNA were excluded from downstream analysis. UMAP cluster annotation was performed manually based on cluster gene expression with the help of the PanglaoDB human datasets and the CellTypist immune cell encyclopedia as primary references^52, 53^. Clusters with less than 150 cells and one cluster high ribosomal RNA content were excluded from downstream analysis.

### BCR repertoire analysis

CellRanger output of BCR libraries was exported for downstream analysis. C gene and clonotype classification was based on the *enclone* output as part of the CellRanger workflow. Violin plots, bar graphs, and heat maps were generated in GraphPad Prism (v9.5.1) along with associated significance testing. Chord diagram for C-gene usage and cluster allocation was generated using TCR_Explore (v1.0)^54^. Clonotypes with n ≥ 4 were analyzed for C gene usage and B cell compartment based on cluster localization in the UMAP. Plasma (C8, C20), germinal center (GC) (C6, C9, C11, C14) or other (C0, C1, C3, C17, C19, C22) groupings were assigned. Filtered BCR contig.fasta files from the CellRanger output were used for V(D)J assignment using IMGT/HighV- QUEST^55, 56^. Germline reconstruction was performed using the *CreateGermlines* function within the Dowser package (v2.1.0)^57^. Mutational frequencies were calculated using the *observedMutations* function as part of the SHazaM R package (v1.2.0)^58^.

### TCR repertoire analysis

CellRanger output of TCR libraries was exported for downstream analysis. TCRs were annotated into CD8+ versus CD4+ based on their expression of *CD8A/B* or *CD4*. In the absence of co-receptor gene expression, TCRs were annotated on the basis of the UMAP cluster they mapped to. TCRs that were unable to be annotated were excluded at this stage. TCRs missing alpha chain sequences or CDR3β sequences were also removed. Final TCR repertoires were composed of paired TCR sequences with CD4 or CD8 annotation. Clonal counts were defined as TCRs with identical Vα-Jα-CDR3α and Vβ-Jβ-CDR3β pairs based on amino acid sequence. If expanded clones were expressed in multiple clusters, they were assigned to their most popular cluster. Pie charts and bar graphs, and associated significance testing were completed using GraphPad Prism (v9.5.1). Chord diagrams were generated using TCR_Explore (v1.0)^54^. Motif alignment was also performed in TCR_Explore using MUSCLE (v3.34.0) and viewed using motifStack (v1.36.1). GLIPH2.0 was used to cluster TCRs of putative similar specificity using default settings and version 2.0 of the reference dataset^19^. Violin plots for specified CD8+ T cell cluster markers were generated using the *FindMarkers* function in Seurat and significance threshold was based on adjusted p-value cutoff ≤ 0.05.

### Collection of appendiceal swabs and DNA extraction for microbiome analysis

Brushes (Cook Medical, #CCB-7-240-3-S) were inserted into the appendiceal lumen upon tissue arrival to Anatomic Pathology in accordance with IRB #21-1241, spun five times, then placed in a sterile tube. A second brush was inserted, spun, and placed in a sterile tube containing 20% glycerol. Both brush samples were immediately placed on dry ice and stored at −80°C. Brush samples were transferred to the Microbiome Metagenomics Facility (MMF) at UChicago for DNA extraction and sequencing. DNA extraction was performed using the QIAmp PowerFecal Pro DNA Kit (Qiagen, #51804).

### 16S rRNA amplicon sequencing

Following DNA extraction, the V4-V5 region of the 16S rRNA genes were PCR-amplified using custom barcoded dual-index primers. Illumina compatible libraries were generated using the Qiagen Q1ASeq 1-step amplicon kit (Qiagen, #180419), and sequencing was performed on the Illumina MiSeq platform using 2×250 paired end reads with a goal of generating 5,000 – 10,000 reads per sample. Raw V4-V5 16S rRNA gene sequence data was demultiplexed by the MMF and fastq files were shared for downstream processing.

### 16S taxonomy assignment and downstream analysis

Quality control for demultiplexed files was performed using dada2 (v1.18.0)^59^. Forward and reverse reads were truncated at 180bp, merged amplicon sequences between 300-360bp were retained, and chimeras were removed using the default consensus method. The resulting amplicon sequence variants (ASVs) was used for taxonomy assignment using the Silva reference database (v138.1) to identify bacteria at the genus level^60^. Additional species level information was added when available on the basis of identical matches between the input ASV and the Silva reference sequence. Downstream analysis was performed using MicrobiomeAnalyst 2.0^61^. Sequencing saturation was assessed using rarefaction curves, and samples were rarefied to the minimum library size of 1,856 reads. Low count reads were filtered out using count ≥ 4, percent prevalence across samples ≥10% to remove low abundance features. This strategy was used for analysis of relative abundance profiles for the top 20 genera, along with alpha and beta diversity. For edgeR and sparCC correlation network analysis, samples were not rarefied, rather total sum scaling (TSS) normalization was applied prior to downstream analysis. EdgeR analysis resulted in 57 significant taxa comparing appendicitis versus control samples using and FDR cutoff ≤ 0.05^62^. For correlation network analysis, the sparCC method was employed, using correlation statistic cutoff of ≥ ±0.3 and p-value of ≤0.05 for inclusion^63^.

### Shotgun metagenomic sequencing

Following DNA extraction, Illumina compatible libraries were generated using the QIAseq FX Library Kit (Qiagen, #180477). Sequencing runs were performed on the Illumina NextSeq1000 platform in the MMF using the 2×150 paired ends reads cassette. Shallow shotgun sequencing was performed with a goal of generating 2-3 million reads per sample.

### Identification of possible microbial peptides using metagenomic sequencing data

Raw reads had adaptors trimmed followed by quality control analysis by Trimmomatic (v.0.39)^64^. Host genome was identified and removed using KneadData (v0.7.10), and contigs were assembled using MEGAHIT (v1.2.9)^65^. Gene calling was performed using Prodigal (v2.6.3)^66^. Functional annotation was performed with eggNOG-mapper (v2.1.12)^67, 68^. Results were annotated as either gram positive or gram negative based on taxonomy predicted from eggNOG-mapper output and input into PSORTb (v3.0.3) to predict cellular localization^69^. Sequences with predicted cytoplasmic protein localization were excluded from downstream analysis, and those with predicted extracellular, cell wall, or outer membrane expression based on PSORTb analysis or eggNOG-mapper function were prioritized for peptide identification. Prioritized sequences were used as input for NetMHCIIpan (v4.3)^70^. To select HLA alleles for inclusion, The Allele Frequency Net Database was referenced, where the top three most frequent alleles for each locus based on

U.S. Caucasian and African-American populations were included to better represent our patient population^71^.

