## Supplemental Figures 1-4 for "Germinal center BCR maturation in appendicitis reveals a role for antigen-specific adaptive immune responses during disease"

**Supplemental Figure Legends**

**Figure S1. Significant GSEA pathways from spatial transcriptomic analysis of Follicular B and T cell regions in appendicitis.**

**(A)** From follicular B cell (FB) segments, significant GSEA pathways as determined using an FWER p-value cutoff of ≤0.05. Pathways are ordered by normalized enrichment score (NES). Pathways relevant to disease pathology are highlighted according to the figure legend.

**(B)** From follicle-adjacent T cell (FT) segments, significant GSEA pathways as determined using an FWER p-value cutoff of ≤0.05. Pathways are ordered by normalized enrichment score (NES). Pathways relevant to disease pathology are highlighted according to the figure legend.

**Figure S2: Summary bubble plots for additional phenotypic genes in T and B single cell clusters.**

**(A)** Bubble plot showing expression of key genes across T cell clusters. Genes are shown the on the y axis, with clusters shown on the x axis. Bubble size corresponds to percent of cells within the cluster expressing that gene and are colored by mean gene expression.

**(B)** Bubble plot showing expression of key genes across B cell clusters. Genes are shown the on the y axis, with clusters shown on the x axis. Bubble size corresponds to percent of cells within the cluster expressing that gene and are colored by mean gene expression.

**Figure S3. Comparison of Resting CD8+ versus Effector CD8+ TCR repertoires in appendicitis.**

**(A)** Frequency of Vβ gene usage in resting CD8+ (C10, brown) versus effector CD8+ (C5, pink) appendicitis TCRs. Each dot represents one donor. All comparisons were non-significant by multiple T tests with Holm-Šídák multiple comparison correction.

**(B)** CDR3β length comparison between C5 and C10 CD8+ T cell populations. Bars show percentage of all TCRs with that length. Differences in length distribution were non-significant by T test.

**(C)** Chord diagrams showing Vβ – Jβ gene pairings in CD8+ T cell populations.

**(D)** Chord diagrams showing Vα – Vβ gene pairings in CD8+ T cell populations.

**Figure S4. Comparison of Resting CD4+ versus Tfh CD4+ TCR repertoires in appendicitis.**

**(A)** Frequency of Vβ gene usage in resting CD4+ (C2, green) versus Tfh CD4+ (C12, gold) appendicitis TCRs. Each dot represents one donor. All comparisons were non-significant by multiple T tests with Holm-Šídák multiple comparison correction.

**(B)** CDR3β length comparison between C2 and C12 CD4+ T cell populations. Bars show percentage of all TCRs with that length. Differences in length distribution were non-significant by T test.

**(C)** Chord diagrams showing Vβ – Jβ gene pairings in CD4+ T cell populations.

**(D)** Chord diagrams showing Vα – Vβ gene pairings in CD4+ T cell populations.

### Figure S1.

**A.**

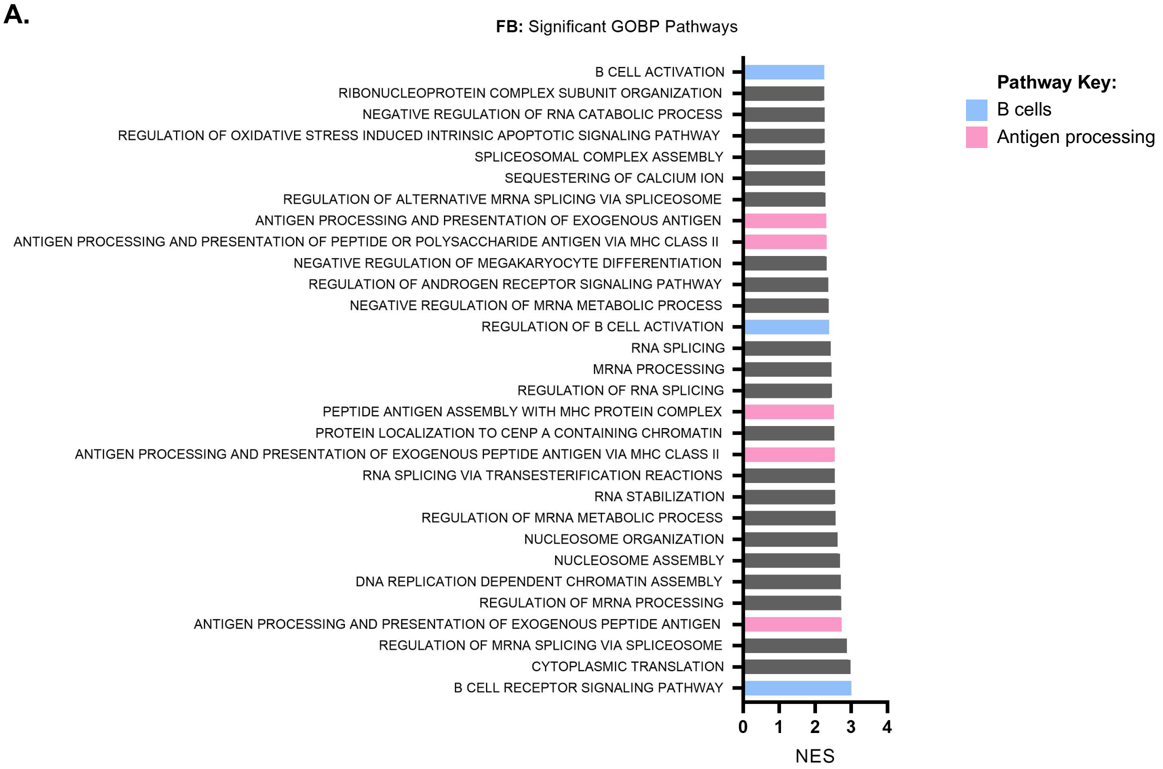
**FB:** Significant GOBP Pathways
B CELL ACTIVATION
RIBONUCLEOPROTEIN COMPLEX SUBUNIT ORGANIZATION
NEGATIVE REGULATION OF RNA CATABOLIC PROCESS
REGULATION OF OXIDATIVE STRESS INDUCED INTRINSIC APOPTOTIC SIGNALING PATHWAY
SPLICEOSOMAL COMPLEX ASSEMBLY
SEQUESTERING OF CALCIUM ION
REGULATION OF ALTERNATIVE MRNA SPLICING VIA SPLICEOSOME
ANTIGEN PROCESSING AND PRESENTATION OF EXOGENOUS ANTIGEN
ANTIGEN PROCESSING AND PRESENTATION OF PEPTIDE OR POLYSACCHARIDE ANTIGEN VIA MHC CLASS II
NEGATIVE REGULATION OF MEGAKARYOCYTE DIFFERENTIATION
REGULATION OF ANDROGEN RECEPTOR SIGNALING PATHWAY
NEGATIVE REGULATION OF MRNA METABOLIC PROCESS
REGULATION OF B CELL ACTIVATION
RNA SPLICING
MRNA PROCESSING
REGULATION OF RNA SPLICING
PEPTIDE ANTIGEN ASSEMBLY WITH MHC PROTEIN COMPLEX
PROTEIN LOCALIZATION TO CENP A CONTAINING CHROMATIN
ANTIGEN PROCESSING AND PRESENTATION OF EXOGENOUS PEPTIDE ANTIGEN VIA MHC CLASS II
RNA SPLICING VIA TRANSESTERIFICATION REACTIONS
RNA STABILIZATION
REGULATION OF MRNA METABOLIC PROCESS
NUCLEOSOME ORGANIZATION
NUCLEOSOME ASSEMBLY
DNA REPLICATION DEPENDENT CHROMATIN ASSEMBLY
REGULATION OF MRNA PROCESSING
ANTIGEN PROCESSING AND PRESENTATION OF EXOGENOUS PEPTIDE ANTIGEN
REGULATION OF MRNA SPLICING VIA SPLICEOSOME
CYTOPLASMIC TRANSLATION
B CELL RECEPTOR SIGNALING PATHWAY
0
1
2
3
4
NES
Pathway Key:
B cells
Antigen processing

**B.**

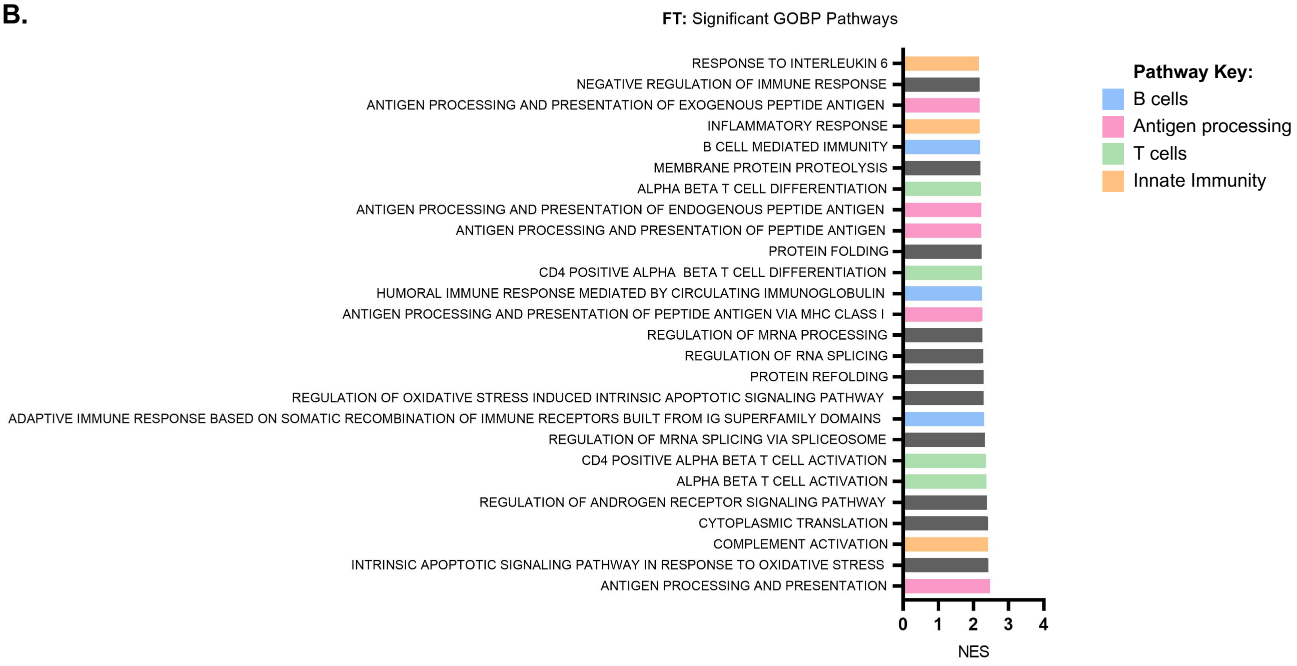
**FT:** Significant GOBP Pathways
RESPONSE TO INTERLEUKIN 6
NEGATIVE REGULATION OF IMMUNE RESPONSE
ANTIGEN PROCESSING AND PRESENTATION OF EXOGENOUS PEPTIDE ANTIGEN
INFLAMMATORY RESPONSE
B CELL MEDIATED IMMUNITY
MEMBRANE PROTEIN PROTEOLYSIS
ALPHA BETA T CELL DIFFERENTIATION
ANTIGEN PROCESSING AND PRESENTATION OF ENDOGENOUS PEPTIDE ANTIGEN
ANTIGEN PROCESSING AND PRESENTATION OF PEPTIDE ANTIGEN
PROTEIN FOLDING
CD4 POSITIVE ALPHA BETA T CELL DIFFERENTIATION
HUMORAL IMMUNE RESPONSE MEDIATED BY CIRCULATING IMMUNOGLOBULIN
ANTIGEN PROCESSING AND PRESENTATION OF PEPTIDE ANTIGEN VIA MHC CLASS I
REGULATION OF MRNA PROCESSING
REGULATION OF RNA SPLICING
PROTEIN REFOLDING
REGULATION OF OXIDATIVE STRESS INDUCED INTRINSIC APOPTOTIC SIGNALING PATHWAY
ADAPTIVE IMMUNE RESPONSE BASED ON SOMATIC RECOMBINATION OF IMMUNE RECEPTORS BUILT FROM IG SUPERFAMILY DOMAINS
REGULATION OF MRNA SPLICING VIA SPLICEOSOME
CD4 POSITIVE ALPHA BETA T CELL ACTIVATION
ALPHA BETA T CELL ACTIVATION
REGULATION OF ANDROGEN RECEPTOR SIGNALING PATHWAY
CYTOPLASMIC TRANSLATION
COMPLEMENT ACTIVATION
INTRINSIC APOPTOTIC SIGNALING PATHWAY IN RESPONSE TO OXIDATIVE STRESS
ANTIGEN PROCESSING AND PRESENTATION
0
1
2
3
4
NES
Pathway Key:
B cells
Antigen processing
T cells
Innate Immunity

### Figure S2.

A.

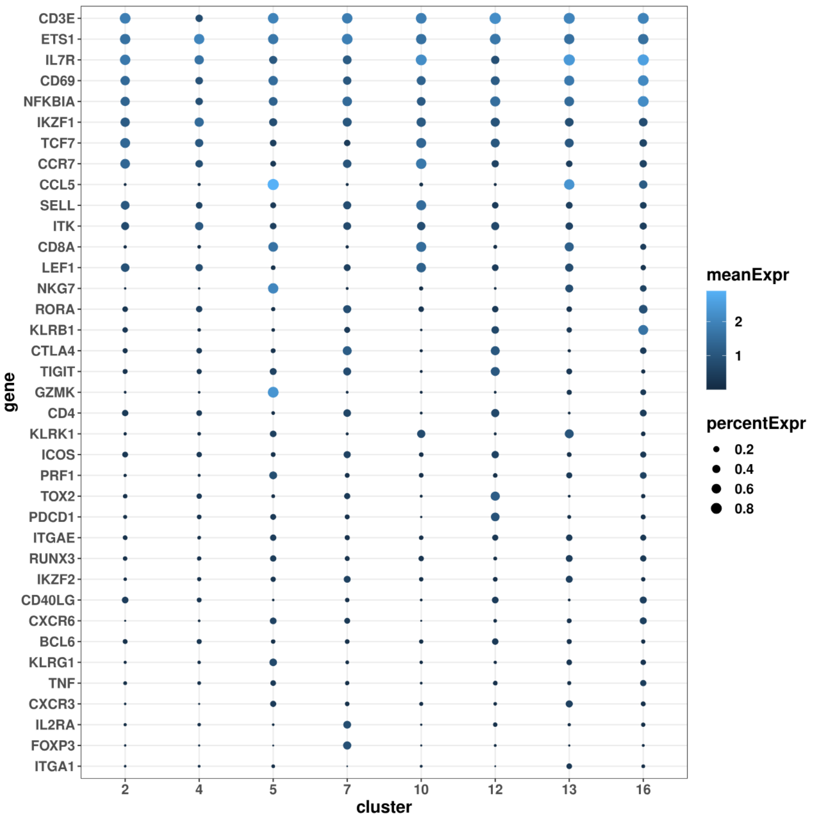

CD3E
ETS1
IL7R
CD69
NFKBIA
IKZF1
TCF7
CCR7
CCL5
SELL
ITK
CD8A
LEF1
NKG7
RORA
KLRB1
CTLA4
TIGIT
GZMK
CD4
KLRK1
ICOS
PRF1
TOX2
PDCD1
ITGAE
RUNX3
IKZF2
CD40LG
CXCR6
BCL6
KLRG1
TNF
CXCR3
IL2RA
FOXP3
ITGA1
gene
2
4
5
7
10
12
13
16
cluster
meanExpr
2
1
percentExpr
0.2
0.4
0.6
0.8


B.

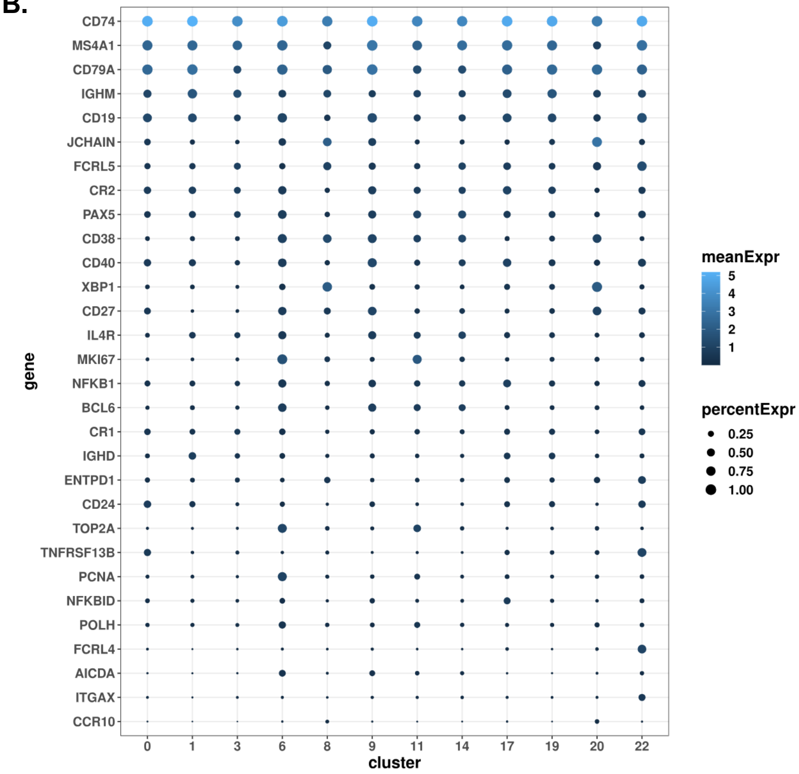

CD74
MS4A1
CD79A
IGHM
CD19
JCHAIN
FCRL5
CR2
PAX5
CD38
CD40
XBP1
CD27
IL4R
MKI67
NFKB1
BCL6
CR1
IGHD
ENTPD1
CD24
TOP2A
TNFRSF13B
PCNA
NFKBID
POLH
FCRL4
AICDA
ITGAX
CCR10
gene
0
1
3
6
8
9
11
14
17
19
20
22
cluster
meanExpr
5
4
3
2
1
percentExpr
0.25
0.50
0.75
1.00

### Figure S3.

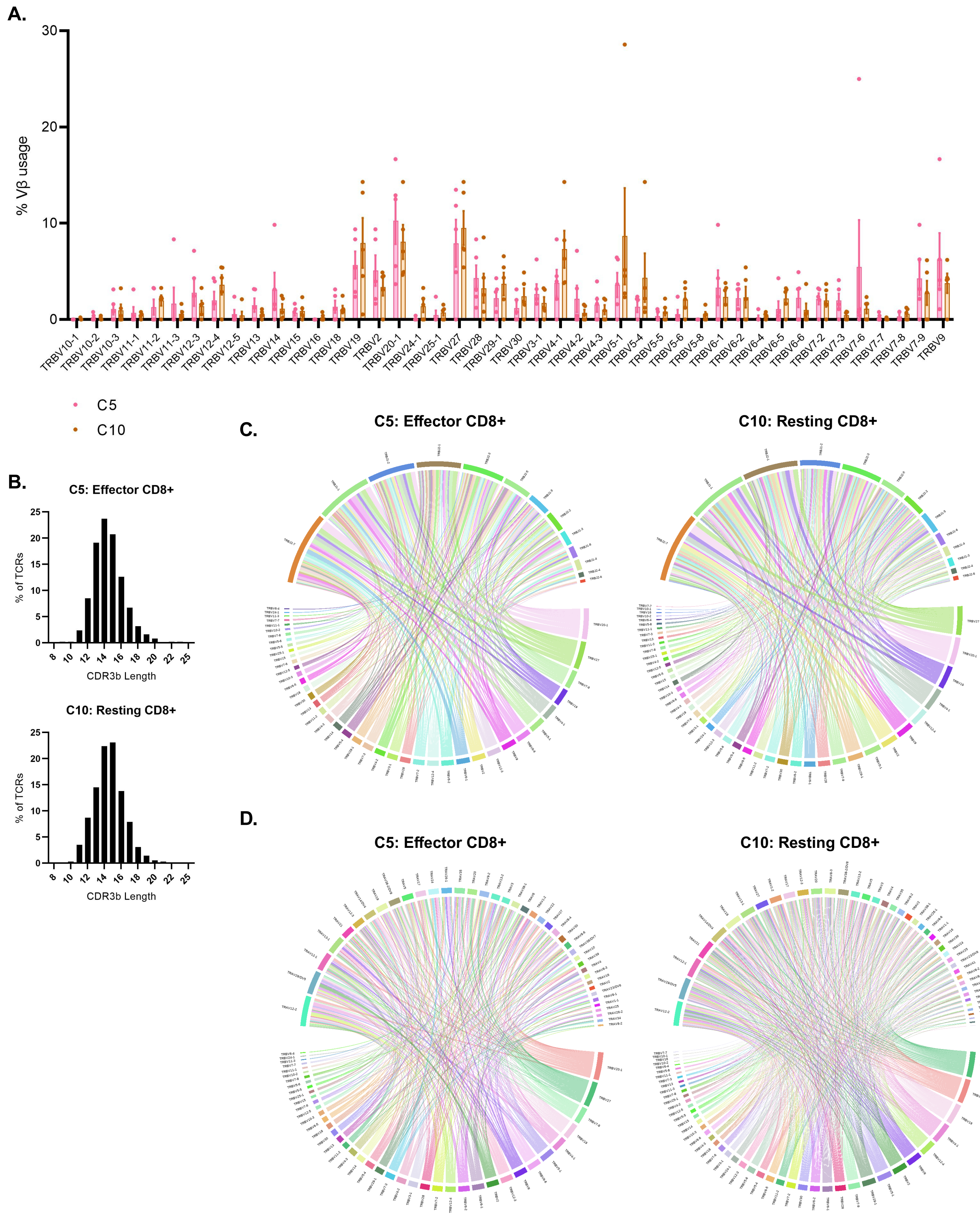
A.
% Vβ usage
30
20
10
0
TRBV10-1
TRBV10-2
TRBV10-3
TRBV11-1
TRBV11-2
TRBV11-3
TRBV12-3
TRBV12-4
TRBV12-5
TRBV13
TRBV14
TRBV15
TRBV16
TRBV18
TRBV19
TRBV2
TRBV20-1
TRBV24-1
TRBV25-1
TRBV27
TRBV28
TRBV29-1
TRBV30
TRBV3-1
TRBV4-1
TRBV4-2
TRBV4-3
TRBV5-1
TRBV5-4
TRBV5-5
TRBV5-6
TRBV5-8
TRBV6-1
TRBV6-2
TRBV6-4
TRBV6-5
TRBV6-6
TRBV7-2
TRBV7-3
TRBV7-6
TRBV7-7
TRBV7-8
TRBV7-9
TRBV9
C5
C10
B.
C5: Effector CD8+
% of TCRs
CDR3b Length
C10: Resting CD8+
% of TCRs
CDR3b Length
C.
C5: Effector CD8+
C10: Resting CD8+
D.
C5: Effector CD8+
C10: Resting CD8+

### Figure S4.

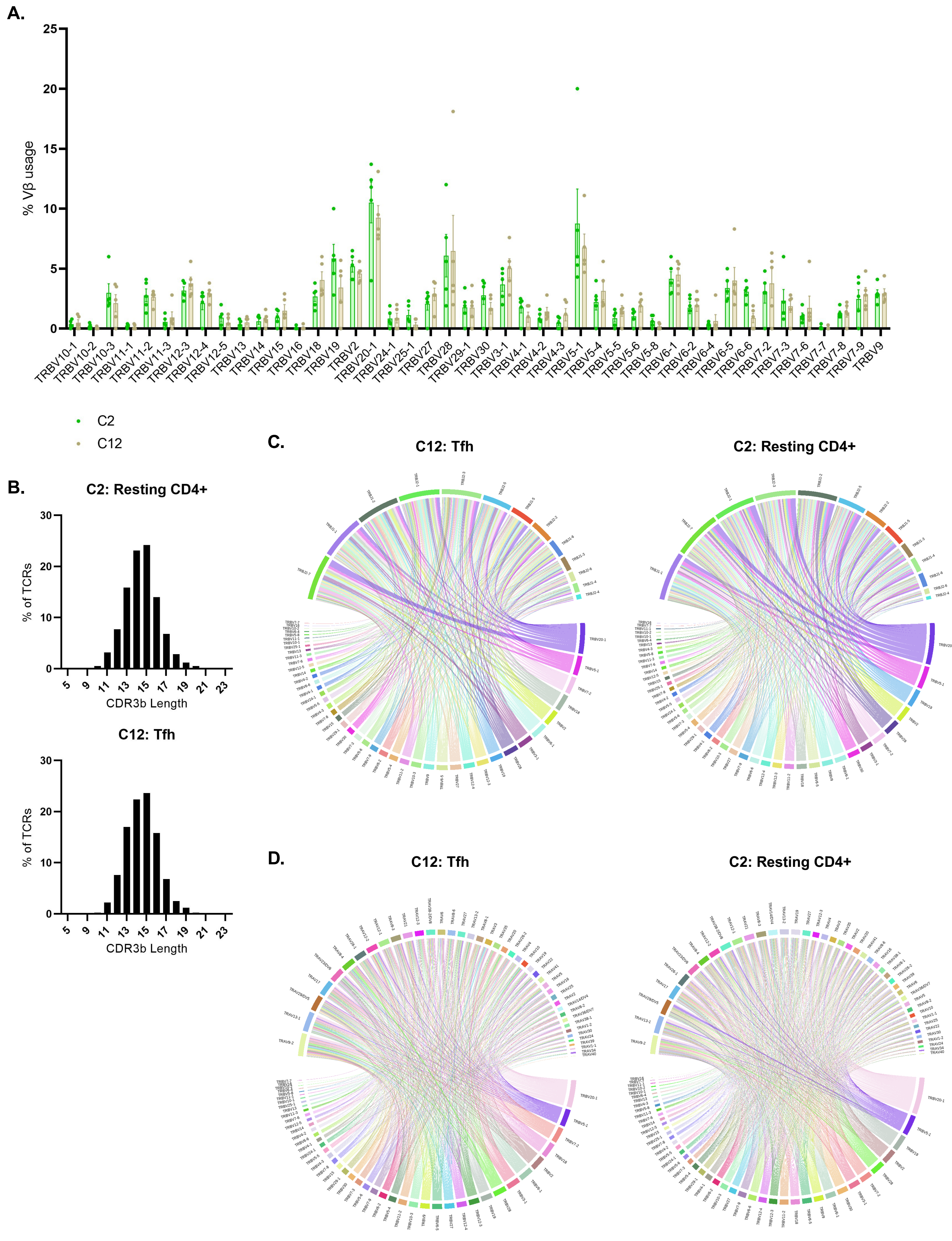
A.
% Vβ usage
25
20
15
10
5
0
TRBV10-1
TRBV10-2
TRBV10-3
TRBV11-1
TRBV11-2
TRBV11-3
TRBV12-3
TRBV12-4
TRBV12-5
TRBV13
TRBV14
TRBV15
TRBV16
TRBV18
TRBV19
TRBV2
TRBV20-1
TRBV24-1
TRBV25-1
TRBV27
TRBV28
TRBV29-1
TRBV30
TRBV3-1
TRBV4-1
TRBV4-2
TRBV4-3
TRBV5-1
TRBV5-4
TRBV5-5
TRBV5-6
TRBV5-8
TRBV6-1
TRBV6-2
TRBV6-4
TRBV6-5
TRBV6-6
TRBV7-2
TRBV7-3
TRBV7-6
TRBV7-7
TRBV7-8
TRBV7-9
TRBV9
C2
C12
B.
C2: Resting CD4+
% of TCRs
30
20
10
0
5
9
11
13
15
17
19
21
23
CDR3b Length
C12: Tfh
% of TCRs
30
20
10
0
5
9
11
13
15
17
19
21
23
CDR3b Length
C.
C12: Tfh
C2: Resting CD4+
D.
C12: Tfh
C2: Resting CD4+
